# Evolutionary divergence in sympatric populations of the fungal pathogen *Alternaria alternata* across wild tomato hosts

**DOI:** 10.1101/2025.10.01.678058

**Authors:** Tamara Schmey, Ben Auxier, Stefan Krebs, Satish Kumar Patneedi, Farooq Ahmad, Michael Habig, Remco Stam

## Abstract

*Alternaria alternata* is a globally distributed fungal pathogen with a broad host range, increasingly affecting both tomato crops and wild tomato relatives. The genomic basis of this ecological breadth in *A. alternata* remains poorly understood. Here we leverage the opportunity of wild pathosystems to study pathogen evolution and diversity beyond agricultural settings. We sampled isolates from wild tomato species across a 2,500 km range in South America, producing highly contiguous genomes, to investigate population structure. Our comparative genomics analyses reveal that *A. alternata* sensu stricto consists of two divergent clades. Strikingly, this divergence is not linked to host species, geography, or habitat type. Transposable elements contribute to variation within clades but do not explain their separation. Although some signs of recombination are present, reproductive mode appears stable across clades. Notably, global reference isolates cluster with one clade, while the other, more diverse clade is only found in wild populations. We hypothesize that these wild populations may act as reservoirs of evolutionary potential. These findings challenge prevailing assumptions about the population structure of necrotrophic pathogens and raise new questions about how genetic divergence can persist without ecological or geographic isolation.

**Media summary (lay abstract):** The fungus *Alternaria alternata* is an emerging threat on tomato and potato crops. In a previous study, we found that it also infects wild tomato plants over a large range in Chile and Peru. Here we sequenced and compared full genomes of the fungi. Surprisingly, they formed two distinct groups that do not reflect different host plant species, geographical locations, or environments. One of those groups is related more closely to global reference samples, while the other group is more diverse. This suggests wild plants may quietly harbour forms of the fungus that could affect crops in the future.

## Background

Wild pathosystems provide essential insights into the ecology and evolution of plant-pathogen interactions, but studies sequencing full genomes of the pathogens remain scarce. Existing work has largely focused on biotrophs like *Microbotryium* [1] and *Uromyces beticola* [2] or hemibiotrophs like *Zymoseptoria* spp. [3] and *Cercospora beticola* [4] leaving necrotrophs comparatively underrepresented. Even though necrotrophs are ubiquitous they have primarily been studied on cultivated hosts, such as for *Sclerotinia sclerotiorum* [5], *Botrytis cinerea* [6], and *Alternaria* species [7].

The genus *Alternaria* comprises filamentous Ascomycete fungi in the family *Pleosporaceae* and includes many saprophytic and pathogenic species across a wide host range [8]. Within *Alternaria*, the taxonomy—particularly of the small-spored species in the *Alternaria alternata* species group—has been frequently revised [9–11]. These small-spored species group are of increasing concern due to their economic and phytosanitary relevance[10]. On tomato crops, they are emerging as dominant pathogens in several regions (e.g. [12–14]), yet they remain understudied compared to large-spored *Alternaria* [15]. Recent work also showed that small-spored *Alternaria* are widespread on wild tomato species [16]. A further example of *Alternaria* on wild hosts is *A. brassicicola* on the coastal halophyte *Cakile maritima* in Australia. Dispersal in this species is partly driven by vertical transmission via ocean currents carrying infected seeds, combined with local spread through both vertical and horizontal transmission [17]. Disease prevalence in these systems is further shaped by host density, environmental factors [18], and pathogen competition. For example, *A. alternata* can also infect *C. maritima*, but is typically outcompeted by *A. brassicicola* in lab settings [17] and additional *Alternaria* species able to infect *C. maritima* are present in Tunisia [19].

Despite the importance of *Alternaria*, the evolutionary dynamics of small-spored *Alternaria* in wild systems remain largely unexplored. In this study, we focus on isolates from eight wild tomato species in Peru and Chile, growing across diverse ecosystems—from the Andes to the Atacama Desert. These hosts are not only valuable models for natural disease resistance [20] but also critical crop wild relatives of cultivated tomato [21]. They exhibit adaptation to both abiotic [22–24] and biotic factors, including variation in resistance to multiple pathogens [25,26]. However, no studies to date have addressed whether *Alternaria* pathogens co-evolve with these wild tomato hosts.

In necrotrophic interactions, the fungi evolve diverse and functionally redundant effectors that induce cell death and sometimes additionally target plant immune functions, while the plant in turn may evolve receptors to recognize these effectors [27]. For *Alternaria*, host specificity is often linked to host specific toxins (HSTs). The biosynthetic gene clusters for the production of the HST reside on accessory chromosomes (ACs), also known as conditionally dispensable chromosomes (CDCs) in *Alternaria* [28]. However, isolates of *A. alternata* can also be pathogenic in the absence of HSTs [11]. *Alternaria* pathogens have a very broad host range, so they might also act as generalists. Its broad host range and global distribution suggest an evolutionary trend toward niche breadth expansion [29].

Transposable elements (TEs) are increasingly recognized for their role in shaping genome plasticity and pathogen evolution [30]. They can inactivate genes, alter gene copy number, and affect gene expression. Genomic regions enriched in TEs often co-localize with pathogenicity-related genes, contributing to the so-called “two-speed genome” architecture in fungi [31]. These dynamic genome compartments, along with horizontal gene transfer (HGT), enable rapid adaptation to new environments or hosts [32]. Recent studies have highlighted the role of giant transposons—termed “starships”—in facilitating HGT, including the transfer of formaldehyde resistance genes across fungal orders [33]. Several genome defense mechanisms exist, among which repeat-induced point mutations (RIP) inactivate repetitive DNA, including TEs, through C-to-T mutations. [34]. Although RIP serves to suppress transposable element activity, it also limits gene duplications that may be beneficial for adaptation [35]. RIP occurs during meiosis hence its signatures in sequences indicates the presence of a sexual cycle. While no sexual stage has been observed in *Alternaria* except for *A. infectoria* [11], accumulating evidence supports the occurrence of recombination and even potential sexual reproduction [7]. This has important implications: clonal reproduction may enhance short-term epidemic spread, while recombination increases long-term adaptive potential in response to environmental and host pressures [36].

In this study, we explore how necrotrophic pathogens evolve in wild systems. We analyse full genomes of 34 small-spored *Alternaria* isolates from wild tomato species collected across diverse South American habitats by Schmey et al. 2023 [16]. We investigate signatures of adaptation to host species, geography, or other selection pressures and assess the potential roles of transposable elements and the reproductive mode in shaping population structure. Using comparative genomics, we provide insights into the distribution, evolutionary dynamics, and potential for adaptation of this globally relevant pathogen.

## Short Methods

Please find extended methods with details about the employed software in the supplementary material.

### Sequencing

The fungal isolates stem from the collection described in [16]. Further information can also be found in supplementary table S1. We extracted DNA using the phenol/chloroform-based method described in Einspanier et al. 2022 [37]. First, we sequenced isolate CS330 on an Oxford Nanopore Technologies (ONT) MinION and obtained illumina short read data. Next, we sequenced 20 isolates on an ONT PromethION using four flowcells and obtained DNBseq short read data for CS046. Afterwards, we performed automated DNA extraction with a KingFisher Flex robot on additional 13 isolates, which we sequenced on an ONT PromethION. Lastly, we obtained DNBseq short read data for eight of those samples. Basecalled ONT reads were filtered with NanoFilt to retain only reads with an average quality score better than Q10.

### Assembly and annotation

We used flye to assemble the long reads and medaka to polish the assemblies. Despite having to correct two assembly artifacts (see extended methods in supplement), this strategy was more reliable than the alternative assemblers and additional scaffolding softwares we compared. For evaluations, we used quast, BUSCO and D-genies. When short read data was available, we polished the respective assembly with three iterations of pilon.

We used the telomeric identification toolkit tidk to verify the telomeric repeat sequence TTAGGG/AACCCT and identify telomeres. For annotation, we used the pipeline funannotate. Repeat annotation and softmasking were conducted with EarlGrey and fed into the funannotate pipeline, as well as biosynthetic gene cluster annotations with AntiSMASH.

### Identification of putative accessory chromosomes

We performed a mapping approach to identify putative accessory chromosomes. To this end, we mapped a concatenated FASTA file with sequence data of all 34 assemblies and nine downloaded *A. alternata* genomes to each of the assemblies and computed coverage statistics using minimap2 and mosdepth. Contigs were classified as putative accessory contigs when they had coverage lower than the mean coverage and at least 100 Kb length. We compared gene density, TE density and gene-wise relative synonymous codon usage (gRSCU) of the putative accessory contigs to the core scaffolds of the respective sample.

We used an *in silico* PCR approach with seqkit amplicon version 2.8.2 to detect the ALT gene for host-specific AAL toxin and a second PKS gene that is used as marker for accessory chromosomes (primer sources see extended methods).

### Phylogeny and k-mer based clustering

We downloaded additional reference genomes (accession numbers see extended methods) and ran BUSCO on them as described above. Then we used the pipeline BUSCO_phylogenomics to produce a supermatrix of concatenated single-copy genes present in all assemblies, which we subsequently used as input for Splitstree, and gene trees that we used as input for ASTRAL. We plotted the ASTRAL tree with information about the isolates in Rstudio. Additionally, we generated a species tree with orthofinder, which we rooted in itol. Mashtree is a k-mer clustering method that provides a tree of the full genome sequences. We ran mashtree on the above-mentioned assemblies and rooted the mashtree in itol. We used phylo.io to compare the orthofinder tree and the mashtree to the ASTRAL tree. Furthermore, we ran mashtree with our assemblies and all available *A. alternata* and *A. tenuissima* assemblies from NCBI, which we rooted to an *A. arborescens* reference and plotted in Rstudio.

### Single Nucleotide Polymorphisms (SNPs)

We mapped the long-read sequencing reads to the Y784-BC03 reference assembly and used the SNP-caller clair3 and filtered the SNPs with bcftools (applied filters see extended methods). After linkage-pruning with plink, we performed a PCA and an ADMIXTURE analysis that we plotted in Rstudio.

We calculated pairwise genetic distances of the samples as Manhattan distances in Rstudio and plotted them as histogram. We used the histogram and the distance matrix to identify clones. Furthermore, we calculated population genetic statistics with Popgenome in Rstudio.

To test for isolation by distance, we retained only biallelic SNPs and calculated genetic distances as Manhattan distances and geographic distances as Haversine distances from the coordinates before correlating the two distance matrices with a mantel test.

### Pangenome construction

To construct a pangenome from the 33 *A. alternata* genomes we used PanTools. After identifying genes that are specific to certain characteristics and phenotypes with the subcommand PanTools gene_classification, we attempted to identify the genes with online Megablast against the NCBI nucleotide database.

### Effector prediction

We ran orthofinder on all proteins and effectorP on the proteins that are predicted to be secreted. Then we identified all orthogroups with at least one predicted effector.

### Selective sweeps

We used RAiSD-AI to detect genomic regions under positive selection in both clades. Then we identified the genes therein and finally performed Gene Ontology (GO) enrichment with BiNGO.

### Transposable elements

To see if repeat content drives the differences in assembly size, we conducted Pearson correlations and Spearman’s correlations on datasets with all samples in total and in both clades before and after outlier removal. To visualize the repeat annotation from EarlGrey, we plotted the HighLevelCount.summary.txt results as stacked bar plots in Rstudio. Additionally, we re-plotted the TE divergence landscapes with fixed y-axis heights.

We used starfish to identify mobile genetic elements termed starships.

### Reproduction

We determined the mating types by conducting a blastn search of the known mating type loci (accession numbers see extended methods) against our assemblies. Then we used binomial tests to check whether mating type ratios deviate from a 50:50 distribution and a chi-squared test to check whether the ratios differ between clades. To investigate repeat induced point mutations, we used the online version of RIPper. We used Welch’s t-test to check whether the percentages of RIP-affected regions differ significantly between clades.

## Results

### Complete genomes do not resolve complete chromosomes

Sequencing yields from ONT PromethION runs varied considerably from 8X to 139X (supplementary table S1). As *Alternaria* has ten core chromosomes [7], the assemblies are at the scaffold level with 13 to 702 scaffolds. One assembly (CS042) was incomplete due to insufficient coverage, all others have BUSCO completeness scores ranging from 97.3% to 99.0%. When ignoring the incomplete assembly, the assembly sizes range from 33.6 Mb to 36.8 Mb. Even when the smallest (RS040) and largest (CS007) are excluded as outliers, the assembly sizes vary from 33.9 Mb to 35.0 Mb. On average (excluding the incomplete assembly), 12049 genes were annotated per assembly. We annotated an average of 36.6 biosynthetic gene clusters (BGCs) per genome, ranging from 33 to 42 per complete assembly (supplementary table S1).

The telomeric identification toolkit tidk confirmed TTAGGG/AACCCT as telomeric repeat sequence. On average, the assemblies showed four to five telomeres, with up to twelve in the best assemblies. Even scaffolds with expected chromosome size rarely had telomeric repeats on both ends. Although some assemblies are too fragmented to allow conclusions about macrosynteny, dotplots reveal that all assemblies are highly syntenic (supplementary figure S1).

There were one to three putative accessory contigs in nine of the 34 assemblies. As expected for accessory chromosomes, the gene density was lower and the TE density was higher in the accessory contigs compared to the core contigs of the same sample (supplementary figure S2A, S2B). However, in only six of the nine cases, the gene-wise relative synonymous codon usage (gRSCU) value of the accessory contigs was significantly different from the value of the core contigs (supplementary figure S2C). There are no biosynthetic gene clusters on the putative ACs except an unidentified one on RS102. None of the 34 assemblies showed genomic evidence for the PKS gene to produce the tomato-specific AAL toxin and the PKS gene to synthesize 6-methylsalicylic acid, which is used as a marker for accessory chromosomes.

### Phylogeny and k-mer clustering reveal lack of apparent specialisation

We built a Splitstree NeighborNet from 5243 BUSCO genes complete and single-copy in all isolates and references genomes and an ASTRAL phylogeny from 6588 gene trees of BUSCO genes in at least four isolates (Figure 1). Both show that all our isolates group with the references for *A. alternata* (Y784-BC03 and Z7) except RS040 which groups with *A. burnsii*. Therefore, we will refer to the remaining 33 except RS040 as *A. alternata* samples. These show a reproducible substructure with two clades, one of which has three subclades and contains all published *A. alternata* references. We confirmed this substructure in a tree from orthofinder and a mashtree from k-mer based clustering of the full genome sequences (supplementary figures. S3 and S4). Plotting the tree with information like host plant species, geographic region and elevation shows that none of these factors explain the observed substructure. Furthermore, we ran mashtree with our isolates and 96 downloaded genomes from NCBI. After rooting to *A. arborescens*, it shows the above-mentioned substructure with all references grouping together in clade 1 (supplementary figure S5). The global reference genomes do not show clear geographical or host specialization, either.

**Figure 1.**
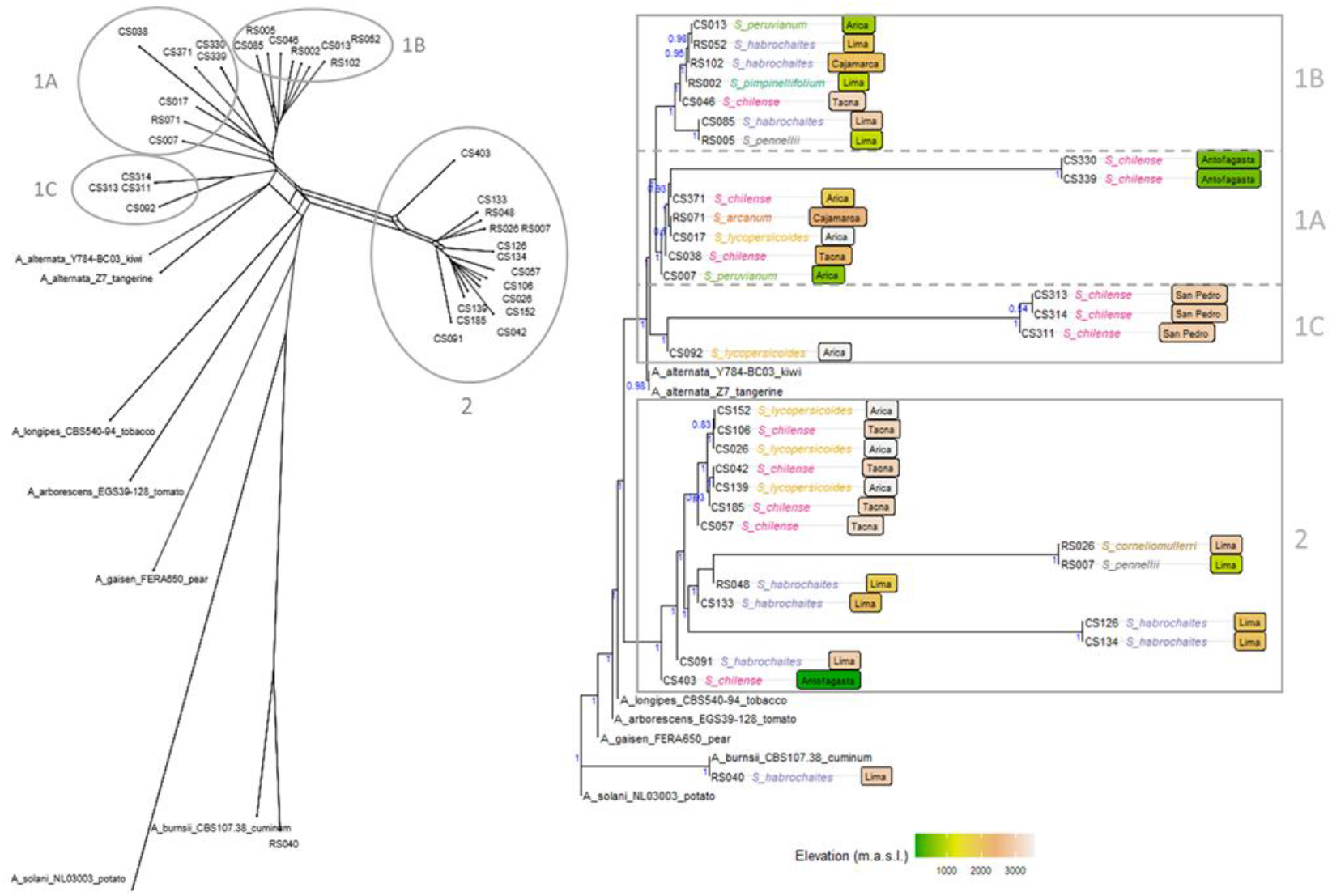
Phylogeny. The NeighborNet from Splitstree (left) is based on a supermatrix of 5243 BUSCO genes. The ASTRAL phylogeny (right) was inferred from 6588 BUSCO gene trees with at least four isolates. We rooted the ASTRAL tree to *A. solani*, and show local posterior probabilities from ASTRAL as blue support values. Coloured tip labels show the *Solanum* host plant species and labels in boxes show the geographic region where the sample was collected with box colour representing the elevation. Clades 1A-C and 2 are consistent across phylogenetic methods.

### Population structure consists of two differentiated clades

The principal component analysis (PCA) clearly shows two groups separating along principal component (PC) 1 (Figure 2A), which corresponds to the phylogenetic groups. The other PCs explained much less of the variation compared to PC1 (Figure 2B). CS403 separates from the other samples along PC2, but remains part of clade 2. As RS040 is not *A. alternata*, it separates from all other samples, which still show the same well-separated groups (supplementary Figure S6).

**Figure 2.**
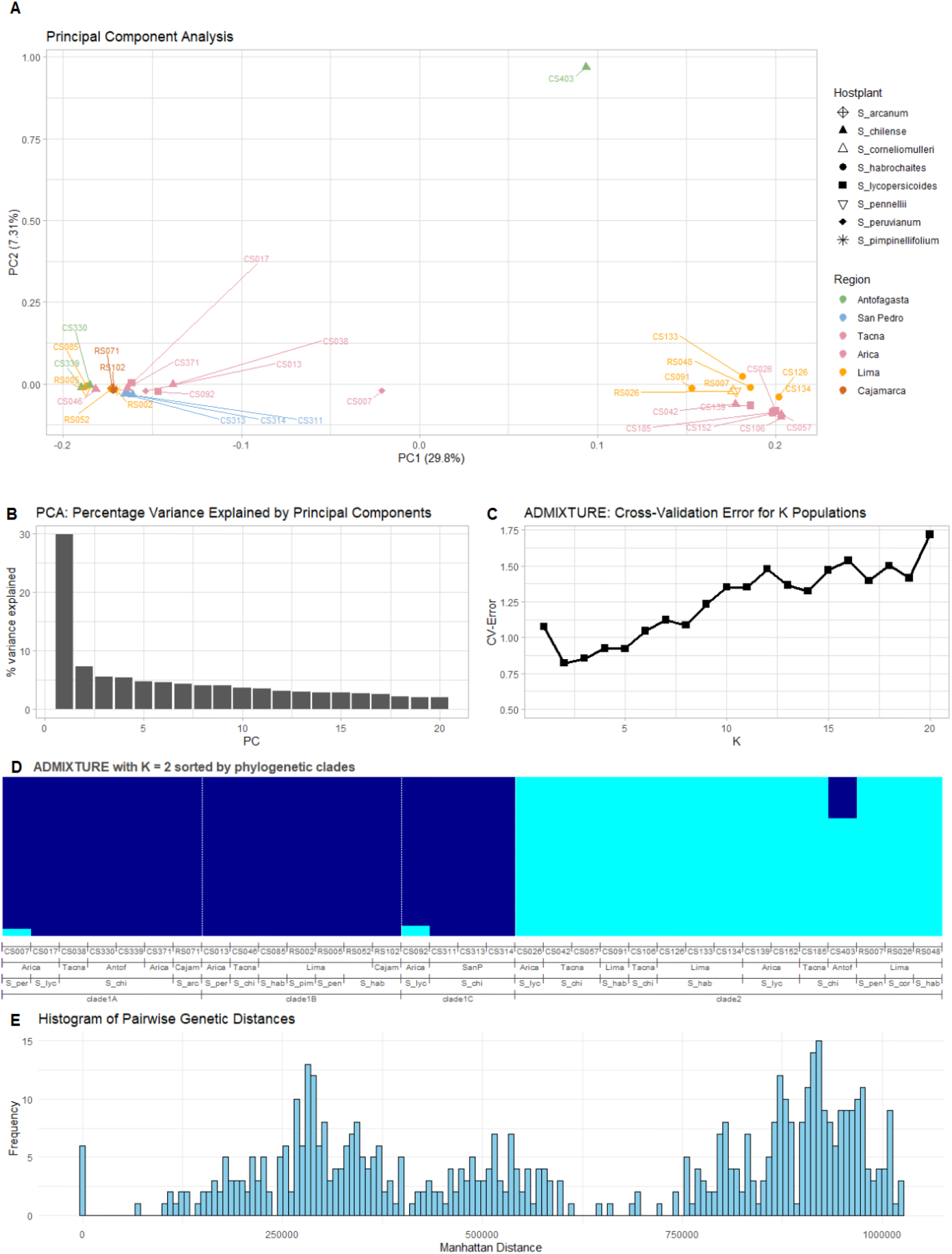
Analyses based on single nucleotide polymorphisms (SNPs) A) Principal component analysis (PCA) of linkage-pruned SNPs to visualize genetic structure among samples. Symbols represent host plant species, and colours indicate collection regions; Tacna and Arica are shown in the same colour to reflect their geographic continuity, as they are both central to the sampling range and separated by a political border. B) The percentage of variance explained by the principal components shown in panel A. C) Cross-validation error of ADMIXTURE analysis for values of K representing the number of inferred subpopulations ranging from 1 to 20. D) ADMIXTURE proportions for K = 2 plotted with samples ordered by phylogenetic clades. E) Histogram of genetic distances between pairs of samples as Manhattan distances.

The cross-validation error for the ADMIXTURE analysis was lowest for K=2 when values up to 20 were tested (Figure 2C). We show that with two groups, the visual representation of the ADMIXTURE analysis confirms the phylogeny and the PCA (Figure 2D).

The histogram of pairwise genetic distances also confirms two distinct subgroups, as there are separate peaks for within-group and between-group comparisons (Figure 2E). The two small peaks with high distances show the comparisons of RS040 to the two groups of all other samples (supplementary figure S7). Six pairwise comparisons have a low distance value and form a histogram bar close to zero, which we define as clones and identify by looking up the lowest values in the distance matrix.

Population genetic diversity statistics like Pi highlight that clade 2 is more diverse than clade 1 (supplementary table S2).

### Selection appears subtle and possibly multifactorial

PanTools considers a gene specific for a phenotype when it is present in all samples that have the phenotype but absent from all other samples. There are no genes specific to any host plant species that occurs more than once in the dataset. No genes are specific to the northern regions, Cajamarca and Lima combined, or to the central regions, Tacna and Arica combined. Fewer collected samples and their higher clonality would cause a bias towards higher numbers of specific genes for southern regions. Summarizing elevations of sampling locations in three bins, there are no genes specific to any elevation ranges. In summary, these results support our hypothesized lack of host, region or elevation specificity. When using the two phylogenetic clades as phenotypes, there are 14 genes specific to clade 1 and 13 genes specific to clade 2. Performing an online Megablast of these specific genes against the NCBI nucleotide database yields mostly hits for uncharacterized and hypothetical proteins from diverse *Alternaria* species.

On average, each isolate has 309 orthogroups with effectors. There are 170 effector-orthogroups shared (core) in all isolates. There are only three effector-orthogroups specific to clade 1 and zero are specific to clade 2.

Selective sweep detection identified 68 and 293 regions (SNP-based window sizes) in Clades 1 and 2, respectively, encompassing 402 and 381 genes. Only 22 genes are under selection in both clades. Most genes were annotated as hypothetical. Functional enrichment of the known genes revealed no significantly enriched GO terms in the clade-specific gene sets. Among shared genes, only the biological process “U6 snRNA 3’-end processing” was significantly enriched. It is involved in splicing, but its adaptive relevance remains unclear.

### The clades differ in Transposable Element composition

Interspersed repeats make up 1.8 % to 4 % of the genome sequences and are positively correlated with assembly size (Spearman’s ρ = 0.755, p < 0.001), especially after removing outliers (ρ = 0.795, p < 0.001). In clade 1, the strong correlation (ρ = 0.771 with outliers, ρ = 0.733 without; both p < 0.01) hints that repeats might drive genome expansion. In contrast, Clade 2 showed weaker and non-significant correlations, particularly after outlier removal (ρ = 0.288, p = 0.32).

On average, repeat contents were higher in clade 2 than clade 1, but variability in clade 2 was high and several samples had repeat contents comparable to clade 1 (figure 3). According to the TE divergence landscapes (supplementary figure S8), signs for recent TE burst occur only in clade 2, but not in all clade 2 samples. In conclusion, while transposable elements likely cause a higher within-clade diversity of clade 2 compared to clade 1, they do not cause the divergence of the two clades.

**Figure 3.**
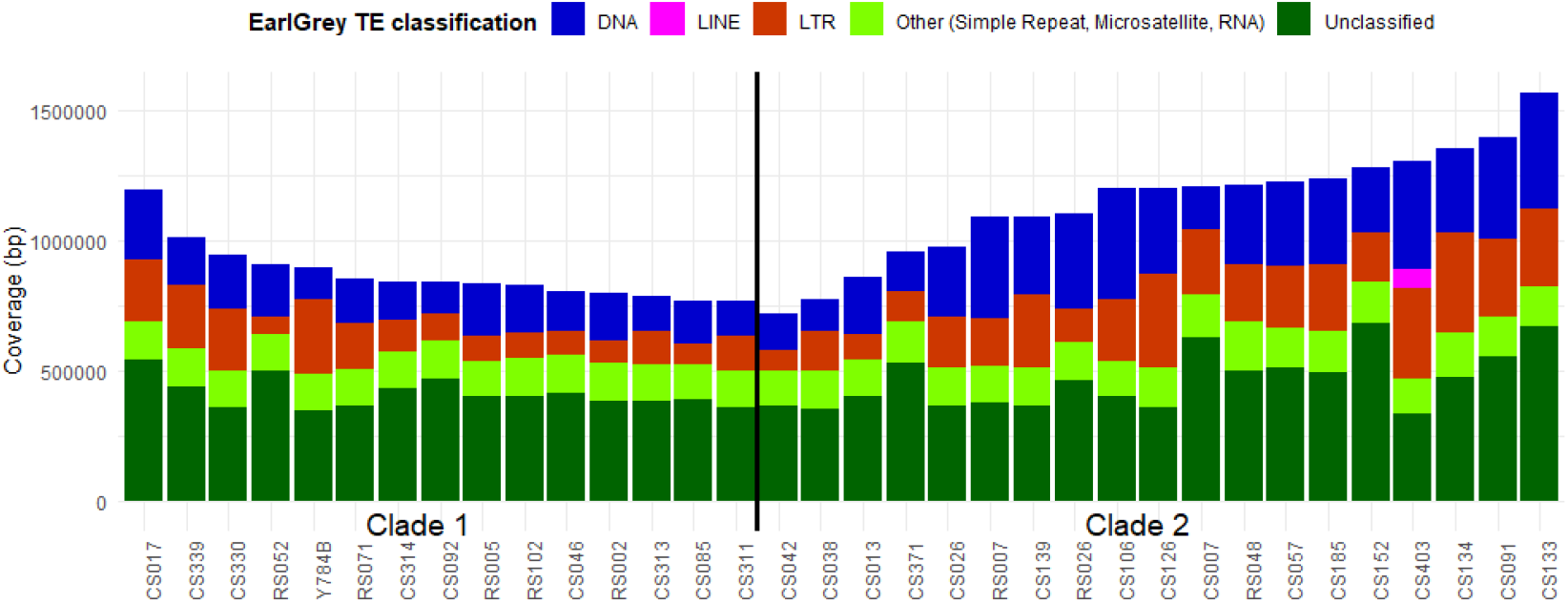
Repetitive sequence content. The **s**tacked bar plot shows the total length of repetitive sequences per sample, sorted in descending order for clade 1 and in ascending order for clade 2. Colours reflect EarlGrey classification.

Across the population, we annotated 24 starship elements, which generally segregated independently from the genome-wide phylogeny (supplementary Figure S9). The most common Region001 in 12 of the genomes had four starship haplotypes but with similar gene content of a predicted histone, and 2-3 proteins of unknown function. Next most common, occurring in seven isolates each, were regions 003 and 004, each with single haplotypes of a few genes with chromatin remodelling and/or transcription factor domains. Other regions carried highly divergent starships, for example Region 5 with two insertions that shared no cargo genes. Surprisingly, there were few secreted proteins in the cargo genes, suggesting only few effector candidates. Other starship elements contain a mix of genes including arsenate reductases and efflux pumps including major facilitatory superfamily (MFS). Most (13) regions also contained predicted proteins with HET domains.

### Cryptic sexuality might contribute to recombination

All isolates have either mating type one (MAT1-1) or two (MAT1-2). The mating type ratios did not significantly deviate from a 50:50 distribution with 18:16 in the total population (binomial test, p = 0.864), 11:7 in Clade 1 (p = 0.481), or 7:8 in Clade 2 (p = 1.000), and the ratios did not differ significantly between clades (Chi-squared test, p = 0.63), suggesting random distribution of mating types across the samples. The mating types occur sympatrically without evident geographic pattern (supplementary figure S10).

Using RIPper, we found that the genomes contain 20 to 47 large RIP affected regions (LRARs) and the percentage of the genomes affected by RIP ranges from 1.4 % to 2.53 % (supplementary table S1). This would indicate an active RIP mechanism in all genomes. Although clade 1 showed a slightly higher average percentage of RIP-affected regions (1.893%) compared to clade 2 (1.771%), this difference was not statistically significant (Welch’s t-test, p = 0.169).

## Discussion

Previously, we found that within *Alternaria* fungi from wild tomato hosts, the small-spored section *Alternaria* was dominant [16]. Building on this study, we sequenced the whole genomes of 34 *Alternaria* isolates from wild tomato hosts in Chile and Peru. Despite the many taxonomic rearrangements of this genus, we confidently identify 33 of the 34 isolates as *A. alternata*. Interestingly, we consistently find that our samples form two distinct subgroups, namely clades 1 and 2. All published *A. alternata* references form a clade that is closely related to our clade 1. However, clade 2 still belongs to the species *A. alternata*, as all three groups form a monophyletic lineage that is sister to the other major species groups identified in [10].

Host specificity of *A. alternata* is usually attributed to host-specific toxins (HSTs), which are encoded by gene clusters located on accessory chromosomes (ACs) [28]. The AC of tomato-infecting *Alternaria* carries the gene for AAL toxin production and a second, unrelated PKS gene called AaMSAS, which are both used as marker genes for ACs [38]. Because the AaMSAS gene is present on the AC of *A. brassicae* without the AAL toxin cluster [39], it could be construed as a more general marker for ACs. Neither of the genes is present in any of our assemblies. As our isolates cause lesions on wild tomato leaves [16], we conclude that AAL toxin is not required for infecting the wild hosts. The absence of AAL toxin is consistent with our species identification, as AAL-producing strains have been reclassified as *A. arborescens* [40].

Whereas host-pathogen co-evolution is typically associated with biotrophic interactions, where tight molecular interplay selects for specificity, necrotrophs also exhibit signatures of local adaptation or population structure linked to host or geography (e.g. [5,41,42]). Interestingly, none of our analyses indicated any grouping by wild tomato host species. Comparison with global samples showed that host plant families were interspersed across the tree, supporting our theory that the core chromosomes exhibit no host-specificity and possibly no adaptation to host plants. More systematic sampling would be required to disentangle true host-associated genomic patterns from confounding effects of shared origin or methodology.

The question whether the wild fungal isolates are capable of infecting domesticated tomato crops was only investigated with one cultivar of tomato and a subset of fungal samples. Additional experiments using more cultivars and entirely different host species are needed to clarify whether the apparent lack of host adaptation reflects true generalism and to assess the potential for host jumps. A study by Peixoto et al. 2021 demonstrated that *Alternaria* from persistent weeds in Brazil could cause disease symptoms on tomatoes and cited additional evidence of cross-infection on various solanaceous hosts [43]. This raises the possibility of the wild system serving as a pathogen genetic reservoir for crop diseases, supported by the higher diversity of the wild clade.

Increased fitness on the original host species or population would be an indicator of host adaptation, which can occur at two levels: through specialization to different host species, or through local adaptation to distinct host populations [44]. In our previous study, we conducted infection assays with a subset of the current samples and showed that pathogenicity was not increased on the original host species. This finding supports our conclusion that the samples do not show host specialization to wild tomato species. The lack of host specialization indicates that the diversity does not come from balancing/fluctuating selection by antagonistic co-evolution with the hosts but is in fact standing variation of the fungal population.

Alternatively, geographic factors could drive differentiation. The correlation of genetic and geographic distance was marginal. This aligns with earlier findings from a multilocus phylogeny of global samples, which also found no association of phylogenetic lineages with host or geography [45]. According to [46], small *A. alternata* conidia could be wind dispersed over 2000 km or even over continents and still be able to germinate. The largest distance between samples from our study is 2150 km, making natural dispersal a likely scenario. This passive and geographically broad dispersal makes fungi more prone to host generalism [29].

Beyond host and geography, no environmental factors seem to drive genomic differentiation at the whole genome and gene presence/absence level. We show that there is no grouping by elevation, indicating that the samples have not locally adapted to this habitat factor. The broad geographic regions represent very distinct climates and habitats, such as the contrast between the humid slopes of the Andes mountains and the Atacama Desert. Given the marked ecological differences between regions, they can serve as a proxy for environmental conditions. Consequently, the influence of more fine-grained habitat factors and climatic variables would be limited. However, it would be an interesting addition to future studies to take advantage of publicly available climate data, especially when more genomes from the sampled regions become available.

Whereas most northern samples were collected one year earlier than the others, the year of collection does not have a meaningful effect on the grouping of the samples. The perennial nature of the host plants also suggests that host availability and population structure would remain stable across consecutive years. Furthermore, previous studies demonstrated that haplotypes of this fungus can persist across growing seasons on annual agricultural hosts, spanning up to five years [47,48].

Both clades exhibit no clear signs of adaptation to known environmental or selective factors and we remain unable to explain their divergence. Most likely, the pathogens evolve in response to selective pressures that remain to be identified. This is supported by selective sweep analysis, which shows only a few genes under selection shared among both clades but many unique genes in each, indicating distinct evolutionary pressures.

Our results suggest that the two clades are sympatric populations infecting multiple shared wild tomato host species. Whereas the generalist necrotroph *Botrytis cinerea* [49] and the anther smut fungus *Microbotryum violaceum* [50] also show divergent populations in sympatry, the populations differentiated on distinct hosts in both cases. Another example of sympatry in anther smut populations is *Microbotryum lychnidis-dioicae* and *M. silenes-dioicae*. However, they not only infect different *Silene* host species but also differentiated in allopatry before being reintroduced [51]. Differentiating sympatric populations of the wilt pathogen *Verticillium dahliae* are associated with pathogenic or endophytic lifestyles [52]. While *A. alternata* is reported as both a pathogen and an endophyte [11], our previous study proved that members of both clades are pathogens causing infection lesions on several wild tomato species [16].

The generally low repeat content in our assemblies is comparable to other studies of *A. alternata* [9,39]. Transposable elements can play a crucial role in adaptive evolution. A study of *Magnaprthe oryzae* found highly clade-specific TE insertions concordant with population divergence [53]. In our study, TE content was higher in clade 2 compared to clade 1 and recent TE bursts were limited to clade 2. However, several members of clade 2 had repeat content and activity as low as clade 1. In conclusion, transposable elements contribute to the within-clade diversity of clade 2, but do not account for the divergence of the two clades.

*Alternaria alternata* is frequently reported to show more genetic diversity than expected for an asexual pathogen [7]. The near-equal frequencies of both mating-types in our samples and the literature (e.g., [54]) suggest sexual reproduction. We found signatures of RIP, that hint a sexual cycle and impact genome evolution by silencing TEs, duplicated genes, and even neighbouring genes (e.g. [55]). Gene tree discordance and recombination signals suggest genetic exchange [11]. Meng et al. 2015 conclude based on SSR markers that sexual reproduction would be more likely than parasexual reproduction [47]. While cryptic sexuality would explain the observed diversity, other mechanisms for clade differentiation such as vegetative incompatibility remain unexplored and could be addressed in future studies.

McMullan et al. 2025 investigated the beet rust pathogen *Uromyces beticola* on wild and domesticated plants. They found two populations, one with all agricultural and a few wild samples and the second with only wild samples and higher diversity [2]. This structure is comparable to our two subpopulations, one containing all published references, which are mainly from crops, and a second one containing only samples from our wild collection and showing higher diversity levels. *U. beticola* reproduces both sexually and clonally. The population with agricultural samples shows higher clonality, consistent with greater benefits to clonal reproduction in an agricultural environment with less host diversity compared to wild host plants. In contrast, our two clades do not differ in reproductive modes, as mating type ratios and RIP-affected regions show no significant differences.

In conclusion, our results reveal the coexistence of two genetically distinct *A. alternata* populations in the same geographic and host environment, without clear ecological or host-related barriers. This raises intriguing questions about how such diversity is maintained in sympatry. Future work should explore microhabitat partitioning, competition, or epigenetic factors as potential drivers. In addition, targeted experimental infection studies could help determine whether subtle differences in pathogenicity or life history strategies exist between the clades, potentially revealing early stages of ecological specialization or incipient speciation and to what extent the second subclade can act as a genetic diversity reservoir.

## Data availability

Supplementary figures, tables and extended methods can be found in the supplementary materials. The sequencing data and annotated genomes have been uploaded to the European Molecular Biology Laboratory’s European Bioinformatics Institute (EMBL-EBI) under the project accession PRJEB93827. All code used in this study has been deposited in Zenodo (DOI 10.5281/zenodo.15768389).

## Acknowledgements

We are grateful to Gonne Clasen for King Fisher DNA extraction and providing sequence data from his project including coordination with sequencing facilities, Sebastian Ploch and Marco Thines for providing sequencing data for 13 isolates, Andrea Tobian Herreno for help with scripts to analyse SNPs, and Manisha Choudhary for help with data upload.

This project was funded by the German Research Foundation (DFG) - Project Number 403835372; FA was funded by a grant of KiTE (Kiel Training for Excellence). The KiTE project has received funding from the European Union’s Horizon Europe research and innovation programme under the Marie Skłodowska-Curie grant agreement No 101081480.

