## Supplementary_materials for "Evolutionary divergence in sympatric populations of the fungal pathogen *Alternaria alternata* across wild tomato hosts"

#### Contents of Supplementary Materials

##### Supplementary figures

Supplementary Figure S1) Dotplots of genome assemblies

Supplementary Figure S2) Putative accessory chromosomes

Supplementary Figure S3) Comparison of ASTRAL phylogeny and Orthofinder species tree

Supplementary Figure S4) Comparison of ASTRAL phylogeny and Mashtree

Supplementary Figure S5) Mashtree with global *A. alternata* references

Supplementary Figure S6) Principal component analysis including all isolates

Supplementary Figure S7) Pairwise genetic distances of all isolates

Supplementary Figure S8) TE divergence landscapes by clade

Supplementary Figure S9) Starship insertions

Supplementary Figure S10) Map highlighting clades and mating types

##### Supplementary tables

Supplementary table S1) List of isolates with information about sequencing, assembly and results

Supplementary table S2) Population genomic statistics

##### Extended methods

With References for Extended Methods

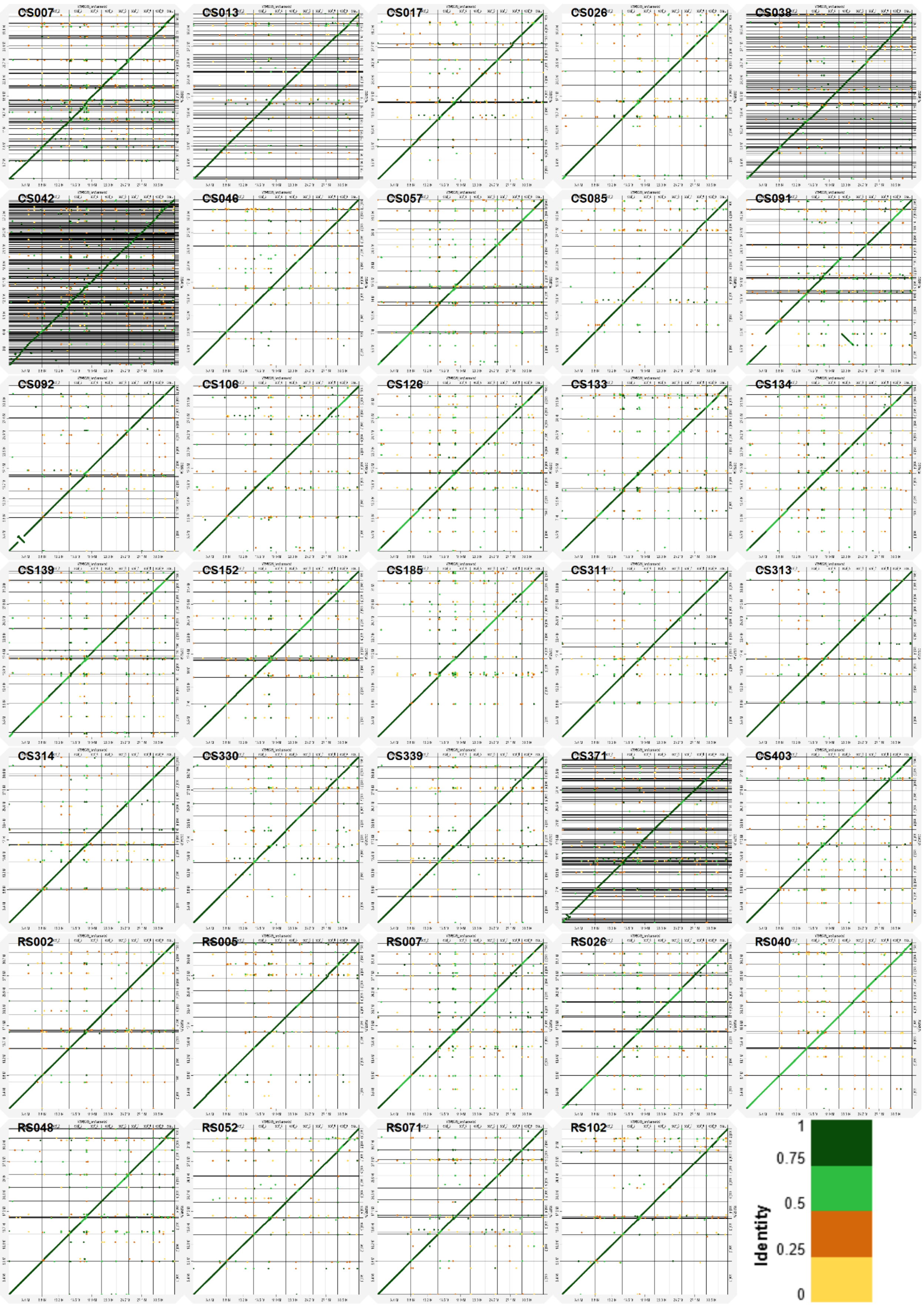


**Supplementary Figure S1) Dotplots of genome assemblies** We mapped all assemblies to the reference genome Y784-BC03 using Minimap2 and visualized alignments with D-GENIES. The reference genome is shown on the x-axis with chromosome boundaries marked by vertical lines; horizontal lines indicate contig boundaries in each assembly. Diagonal lines represent synteny between contigs and the reference. The color for alignment identity indicates sequence similarity.

A

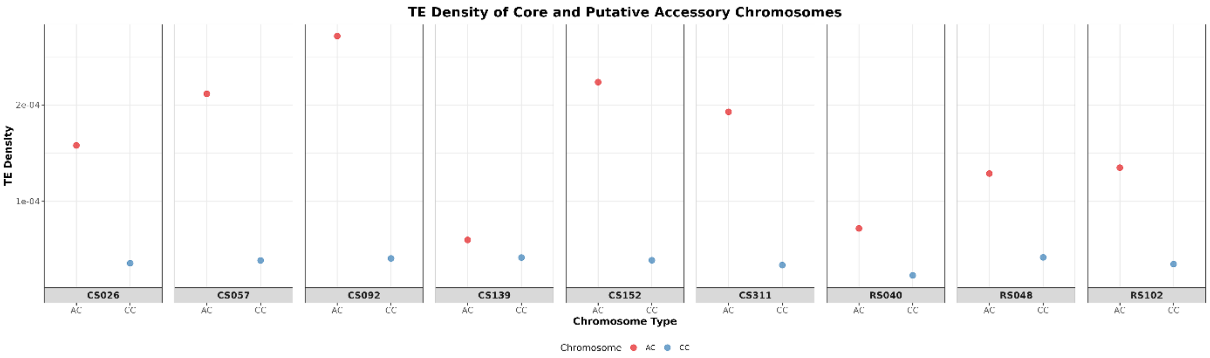


B

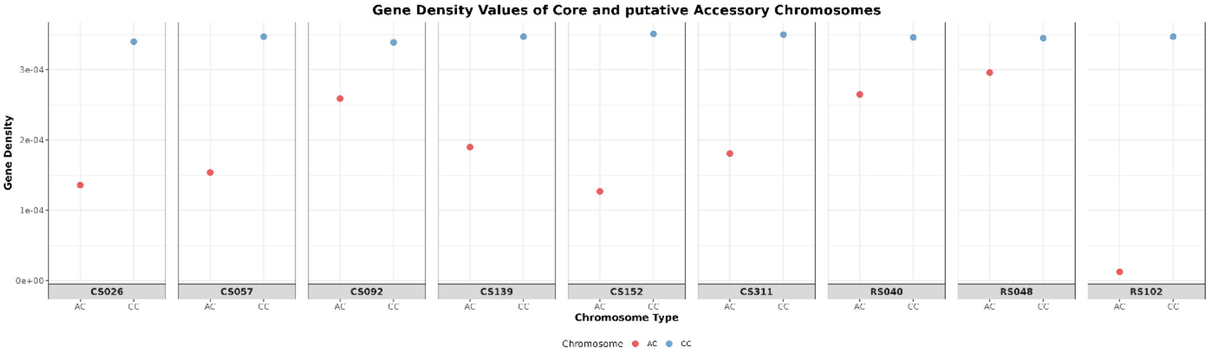


C

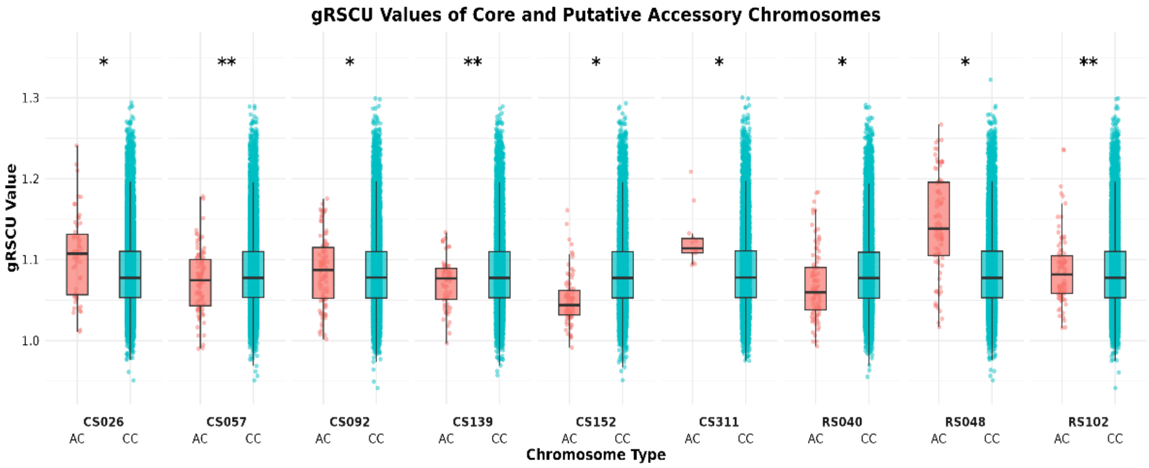


**Supplementary Figure S2) Putative accessory chromosomes** Comparison of putative accessory chromosomes (AC, red/salmon) to the core chromosomes (CC, light blue/teal) of the respective assembly regarding (A) TE density, (B) gene density and (C) gene-wise relative synonymous codon usage (gRSCU) values. Box plots illustrate the distribution of codon usage bias measurements, with statistical significance determined by Wilcoxon rank-sum test. Double asterisks (**) denote isolates exhibiting no significant differences in codon usage patterns between AC and CC contigs (CS057, CS139, RS102), suggesting vertical inheritance of these contigs. Single asterisks (*) indicate isolates with statistically significant codon usage bias differences between putative accessory and core contigs (CS026, CS092, CS152, CS111, RS040, RS048), consistent with horizontal chromosomal acquisition events.

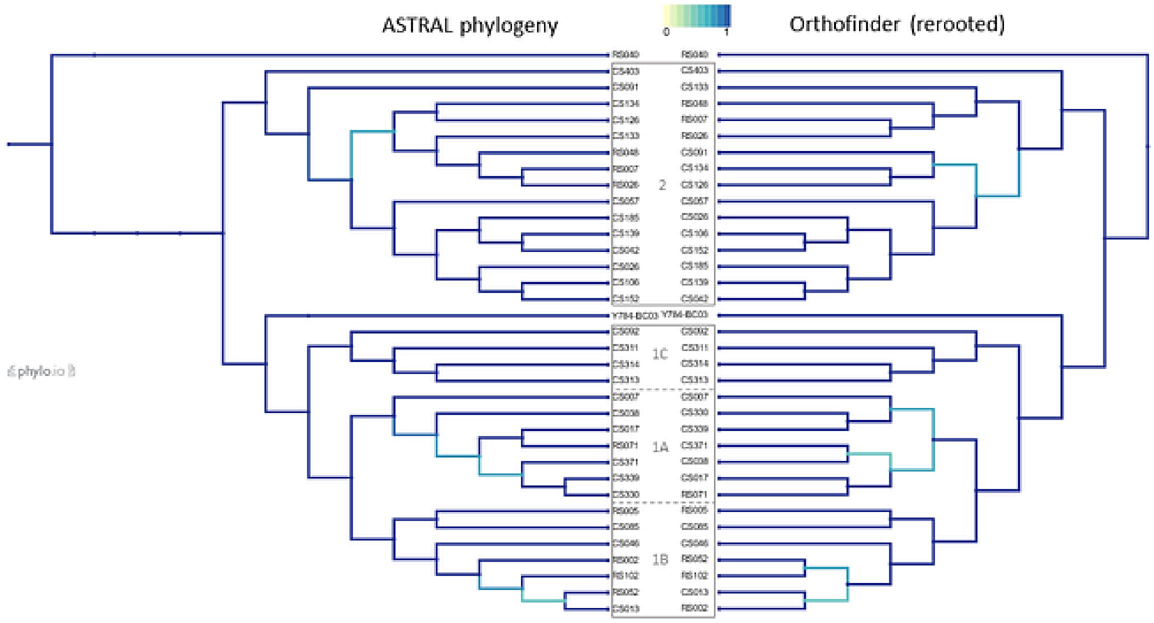


**Supplementary Figure S3) Comparison of ASTRAL phylogeny and Orthofinder species tree** The ASTRAL tree is the ASTRAL tree from figure 1 after pruning references in iTOL. The Orthofinder tree was rooted to RS040 in iTOL. We used Phylo.io to calculate and visualize the tree comparison. Dark blue branches indicate identical nodes (similarity score = 1); lighter colours show lower similarity. Grey boxes in the center mark phylogenetic clades defined in figure 1.

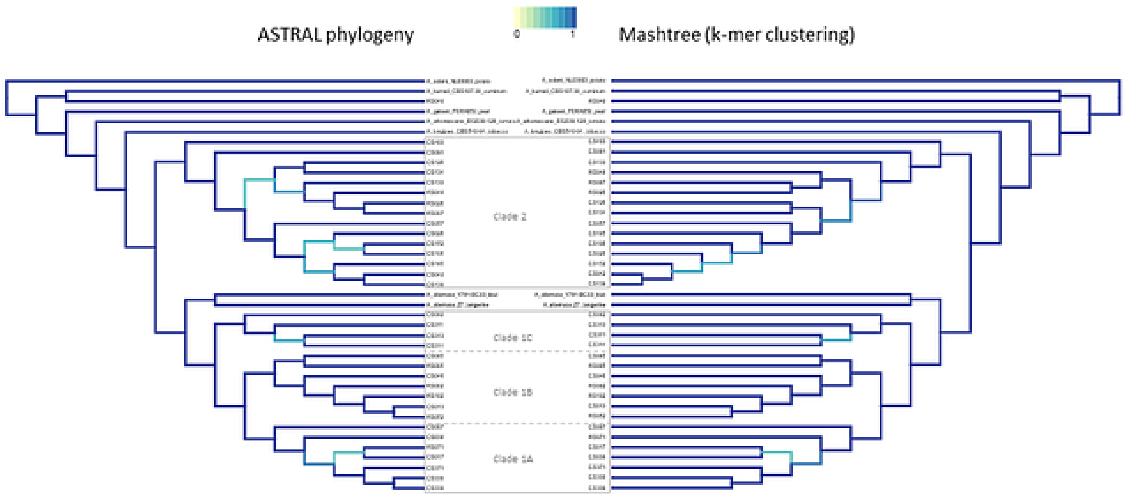


**Supplementary Figure S4) Comparison of ASTRAL phylogeny and Mashtree** The ASTRAL tree is the ASTRAL tree from figure 1. The Mashtree was rooted to *A. solani* in iTOL. As above, we used Phylo.io to calculate and visualize the tree comparison. Dark blue branches indicate identical nodes (similarity score = 1); lighter colours show lower similarity. Grey boxes in the center mark phylogenetic clades defined in figure 1.

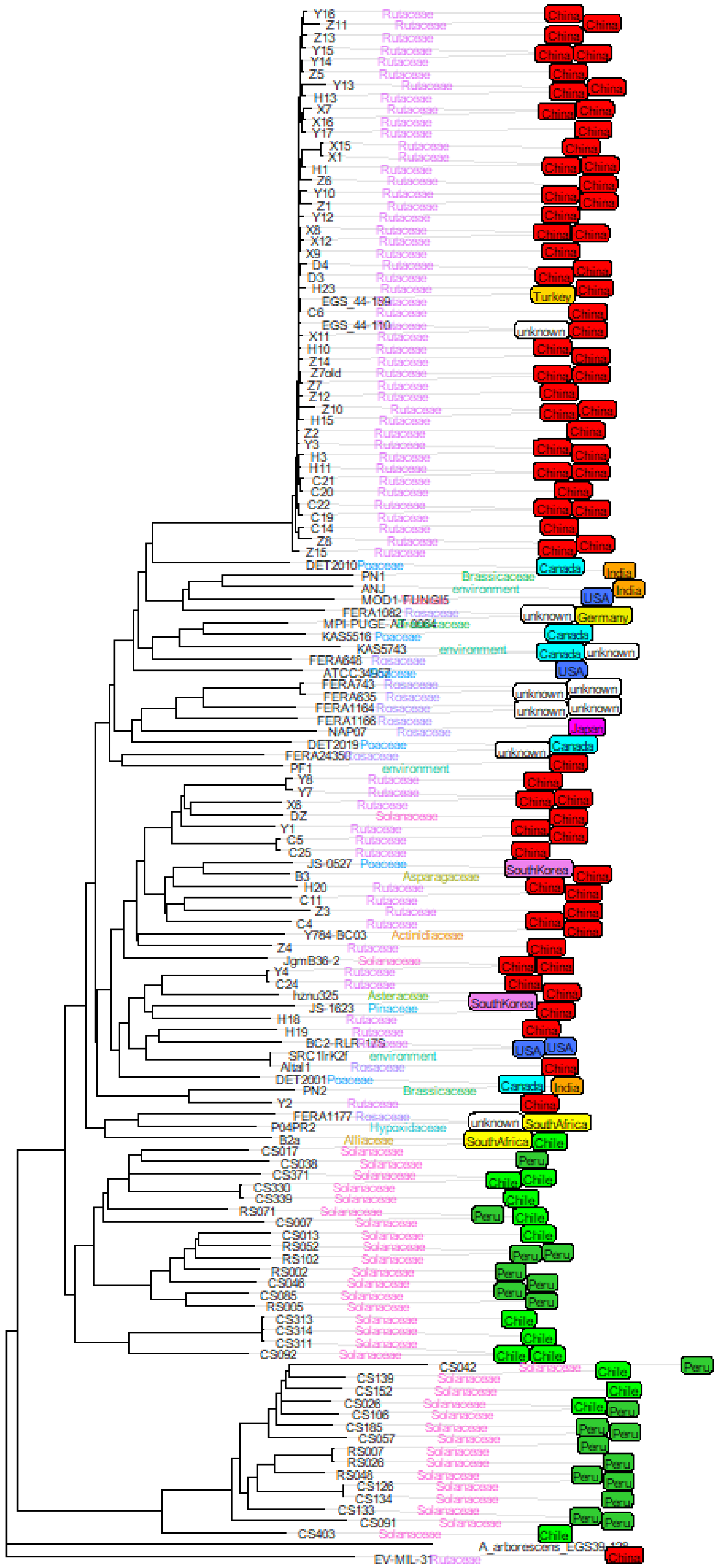

**Supplementary Figure S5) Mashtree with global *A. alternata* references** We constructed a Mashtree using a k-mer–based clustering method, including our 33 *A. alternata* isolates (with green labels for Chile and Peru) and 99 publicly available genomes (89 *A. alternata*, 9 *A. tenuissima*, and 1 *A. arborescens*). We rooted the tree with the *A. arborescens* genome. The black tip labels show isolate names, the coloured tip labels indicate the host plant family, and boxed labels show the country of origin.

A

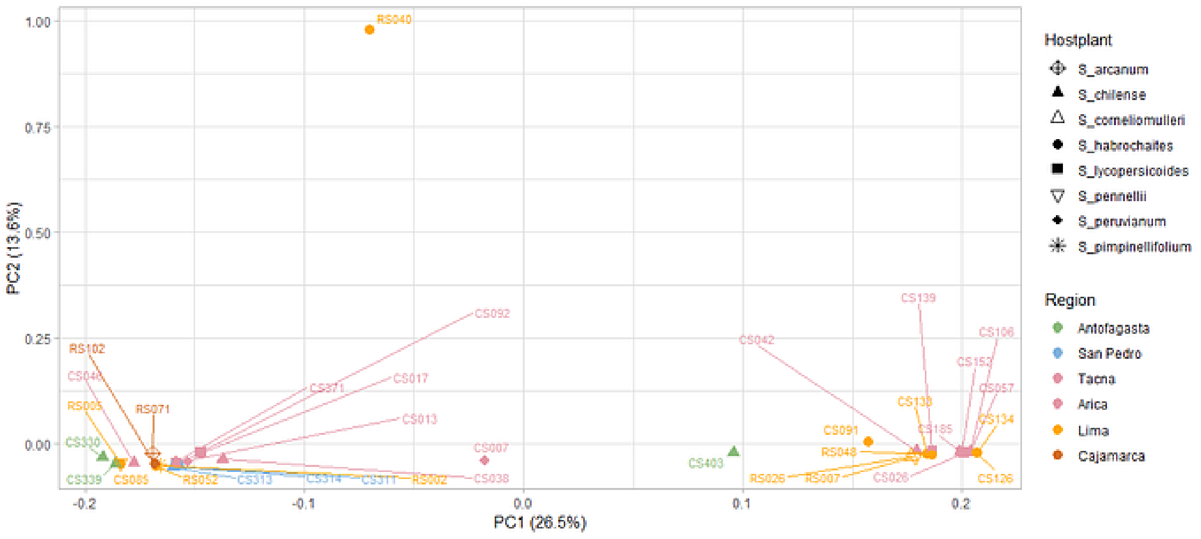


B

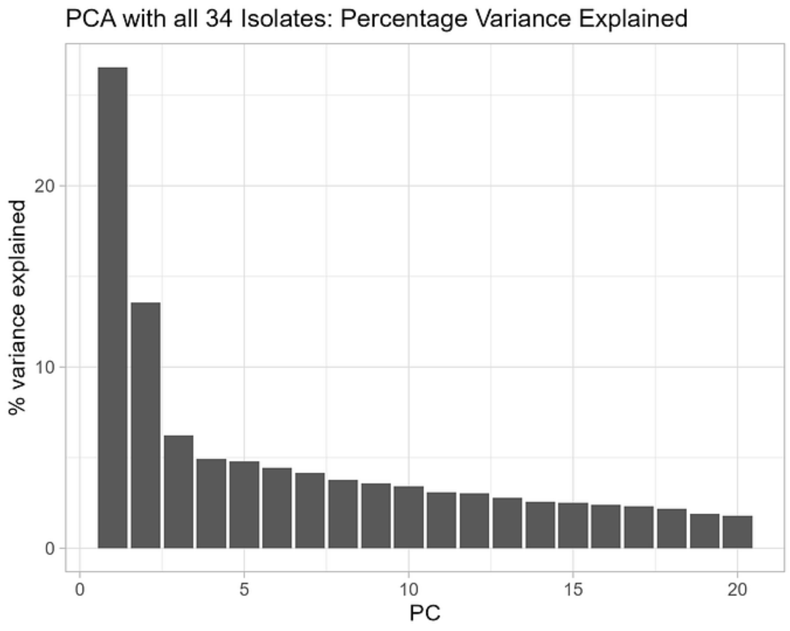


**Supplementary Figure S6) Principal component analysis including all isolates** A) PCA plot constructed as figure 3A, but this version includes RS040, which was excluded from the main figure. B) Percentage of variance explained by the principal components shown in panel A. The plot corresponds to figure 3B but includes RS040.

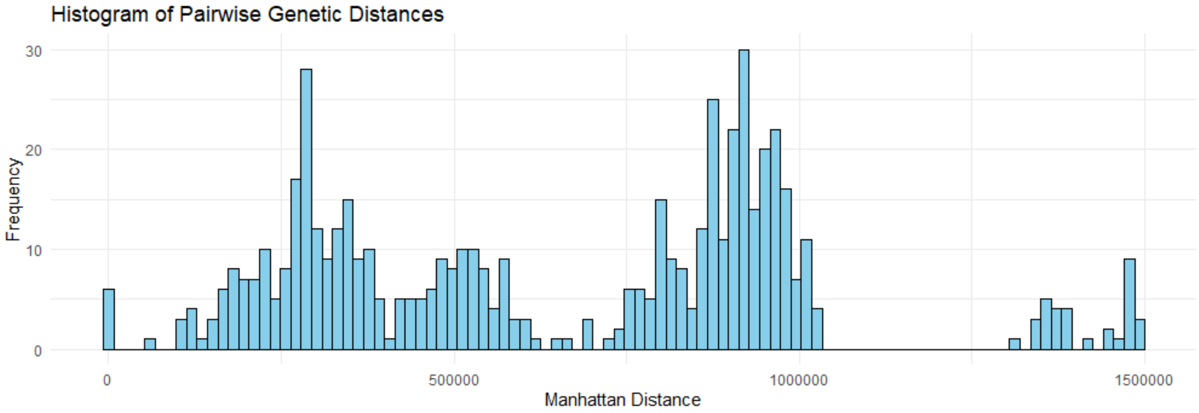


**Supplementary Figure S7) Pairwise genetic distances of all isolates** Histogram of genetic distances between pairs of samples as Manhattan distances. This plot corresponds to figure 3E but includes isolate RS040.

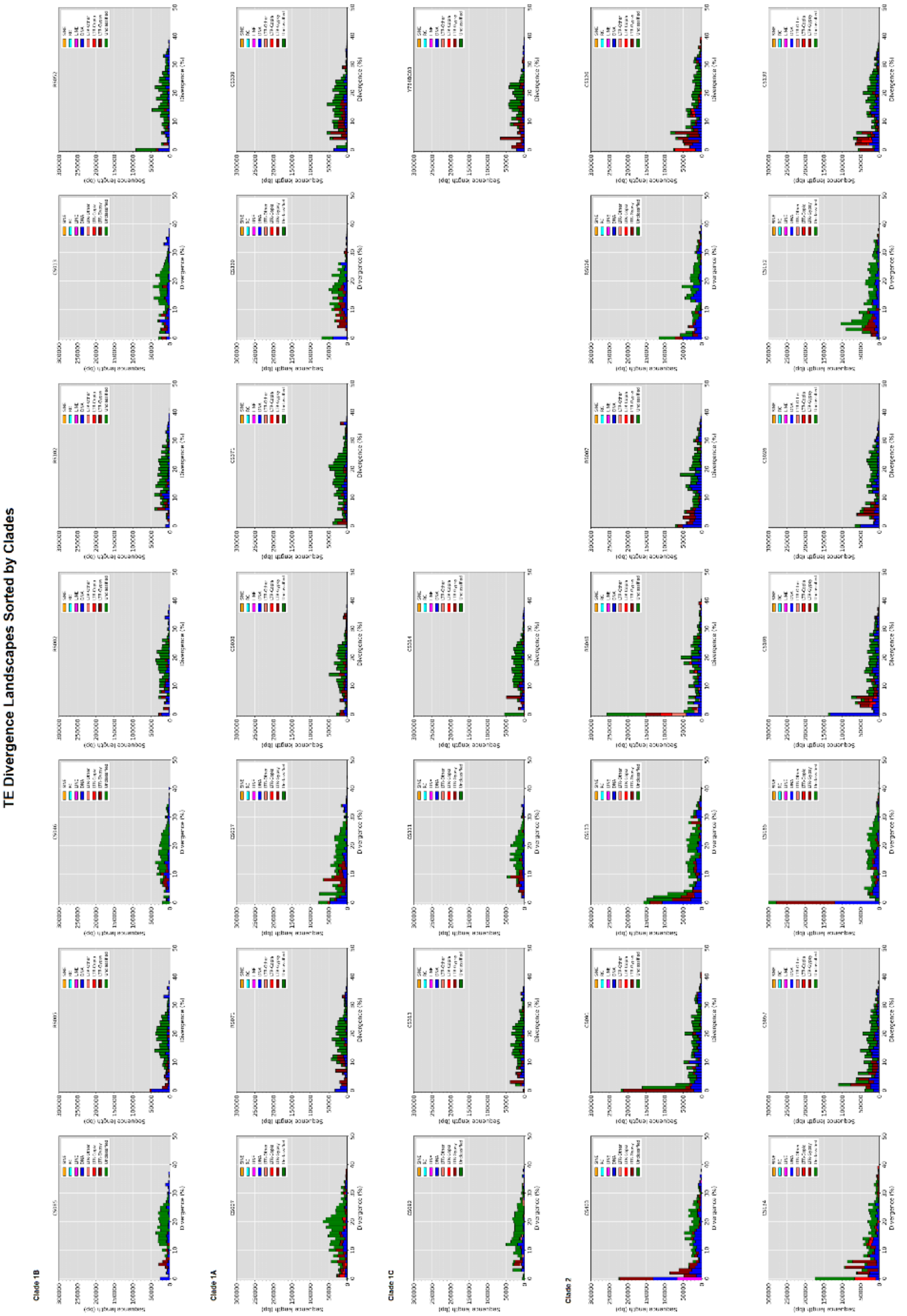


**Supplementary Figure S8) TE divergence landscapes by clade** TE divergence landscapes, generated by EarlGrey, are sorted in rows for phylogenetic clades. The reference genome Y784-BC03 belongs to clade 1 and is depicted behind clade 1C. To make all landscapes comparable, we modified the axis to always show up to 300 kbp, which unfortunately cuts off the first bar for CS185.

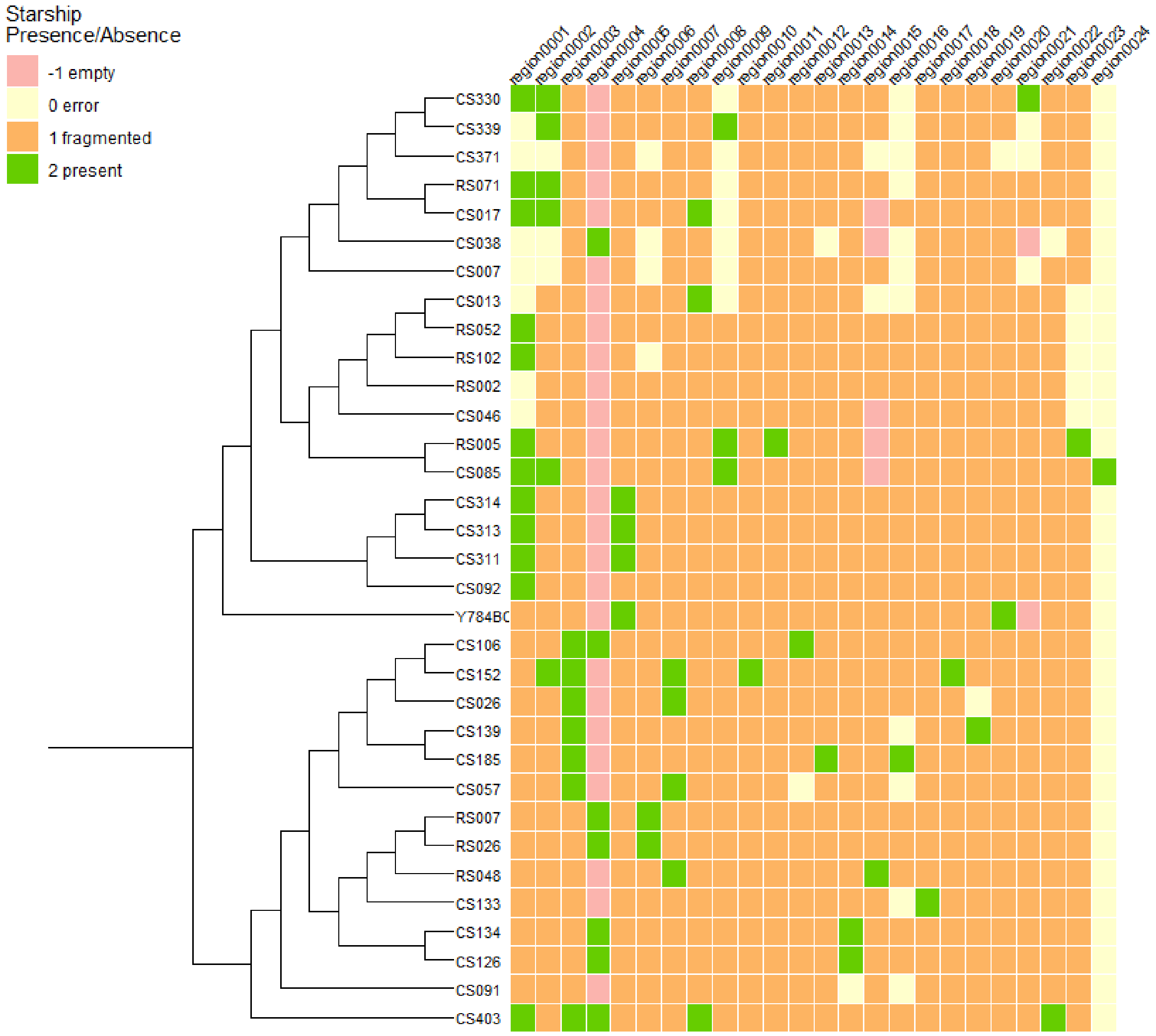


**Supplementary Figure S9) Starship insertions** Starfish determined 24 regions with starship insertions, which are shown in columns. Colours indicate the presence or absence of a starship in this region. Rows representing isolates are sorted by phylogeny, the phylogenetic tree is the ASTRAL tree (see figure 1) after pruning irrelevant rows in itol (see supp. Figure S3).

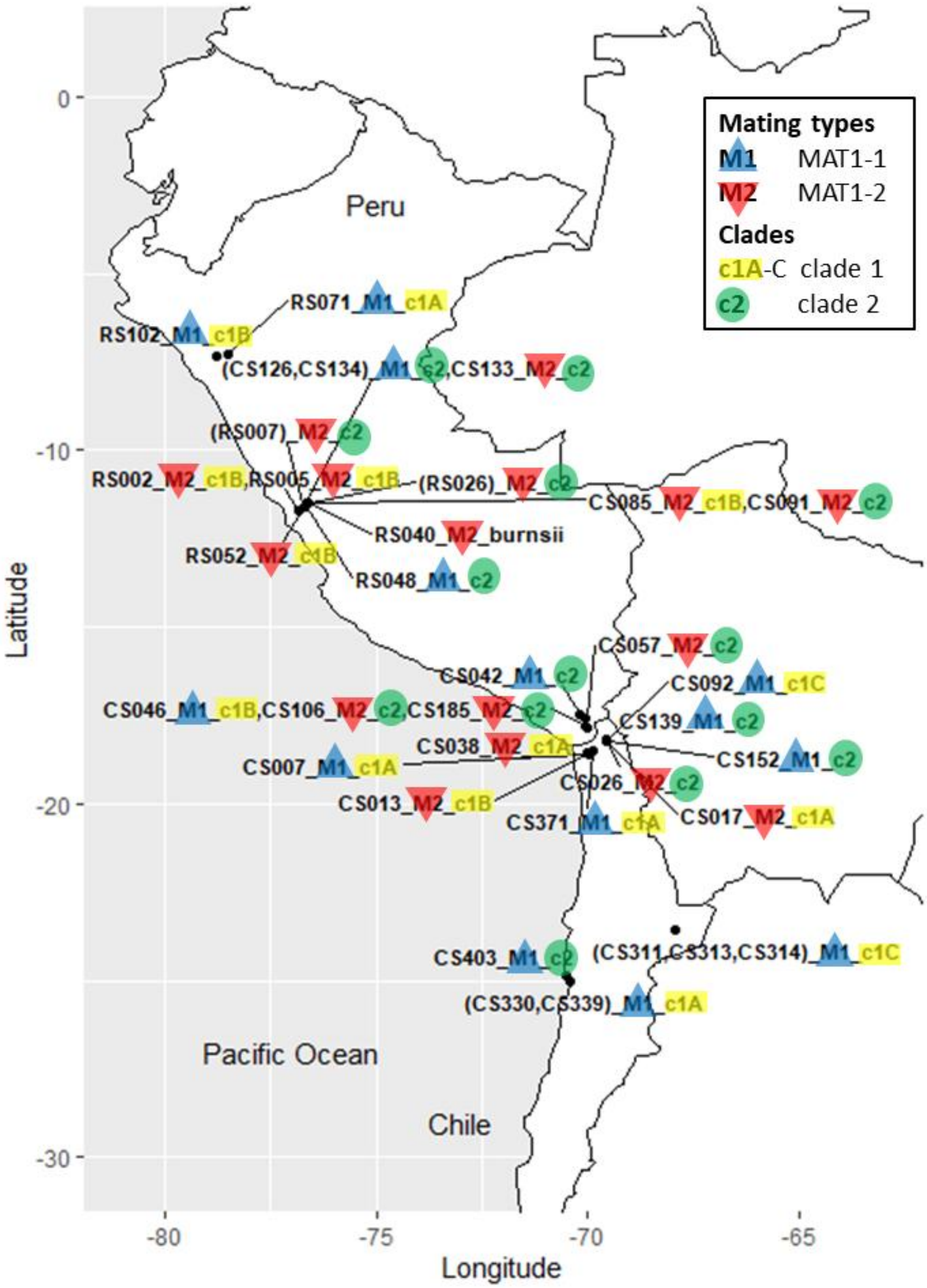


**Supplementary Figure S10) Map highlighting clades and mating types** Samples from the same collection location are displayed in one row, brackets indicate clones. Both mating types MAT1-1 in blue and MAT1-2 in red occur in both clades. Clade 1 is highlighted in yellow and clade 2 in green.

**Supplementary table S1) List of isolates with information about sequencing, assembly and results** ONT batch explanation: 0 for sample CS330 which was sequenced as pilot project on a MinION device, 1 represents the first batch of 20 samples sequenced on a PromethION device and 2 represents the second group consisting of 13 samples. Mean sequencing coverage of the ONT data is reported from the output of the assembler flye. Assembly statistics and GC content were determined with quast.

| **Isolate** | **Hostplant** | **Region** | **Elevation (m)** | **ONT batch** | **Short read polished** | **Mean ONT coverage (flye)** | **Number of contigs** | **Assembly length (bp)** | **N50** | **L90** | **BUSCO scores** |
|---|---|---|---|---|---|---|---|---|---|---|---|
| CS007 | S_peruvianum | Arica | 609 | 1 | no | 43 | 471 | 36773989 | 1073808 | 60 | C:99.0%[S:98.3%,D:0.7%],F:0.3%,M:0.8%,n:6641,E:2.1% |
| CS013 | S_peruvianum | Arica | 779 | 1 | no | 16 | 138 | 33967444 | 653325 | 50 | C:98.6%[S:98.5%,D:0.1%],F:0.4%,M:1.0%,n:6641,E:1.9% |
| CS017 | S_lycopersicoides | Arica | 3571 | 1 | no | 28 | 49 | 34641470 | 2932415 | 11 | C:99.0%[S:98.9%,D:0.1%],F:0.3%,M:0.8%,n:6641,E:1.9% |
| CS026 | S_lycopersicoides | Arica | 3589 | 2 | yes | 50 | 20 | 34177810 | 3107157 | 9 | C:98.9%[S:98.9%,D:0.1%],F:0.3%,M:0.8%,n:6641,E:2.1% |
| CS038 | S_chilense | Tacna | 2221 | 2 | no | 8 | 167 | 33880814 | 359072 | 86 | C:97.7%[S:97.3%,D:0.3%],F:0.4%,M:1.9%,n:6641,E:2.8% |
| CS042 | S_chilense | Tacna | 3078 | 1 | no | 19 | 702 | 30210009 | 66281 | 450 | C:85.3%[S:84.6%,D:0.8%],F:1.0%,M:13.6%,n:6641,E:2.1% |
| CS046 | S_chilense | Tacna | 3091 | 1 | yes | 57 | 42 | 33988710 | 3036740 | 10 | C:99.0%[S:98.9%,D:0.1%],F:0.3%,M:0.7%,n:6641,E:2.0% |
| CS057 | S_chilense | Tacna | 3260 | 1 | no | 108 | 69 | 35000570 | 2820609 | 12 | C:98.9%[S:98.7%,D:0.2%],F:0.3%,M:0.8%,n:6641,E:2.1% |
| CS085 | S_habrochaites | Lima | 2921 | 2 | yes | 139 | 16 | 33888872 | 3051386 | 9 | C:99.0%[S:98.9%,D:0.1%],F:0.3%,M:0.7%,n:6641,E:1.9% |
| CS091 | S_habrochaites | Lima | 2921 | 2 | yes | 15 | 58 | 34712582 | 2384593 | 14 | C:98.9%[S:98.8%,D:0.1%],F:0.3%,M:0.8%,n:6641,E:2.1% |
| CS092 | S_lycopersicoides | Arica | 3533 | 2 | no | 21 | 49 | 34200984 | 2663866 | 12 | C:98.9%[S:98.8%,D:0.1%],F:0.3%,M:0.8%,n:6641,E:1.9% |
| CS106 | S_chilense | Tacna | 3091 | 2 | yes | 60 | 20 | 34511660 | 3123375 | 9 | C:98.9%[S:98.9%,D:0.1%],F:0.3%,M:0.8%,n:6641,E:2.1% |
| CS126 | S_habrochaites | Lima | 2047 | 2 | yes | 79 | 23 | 34491588 | 3123892 | 10 | C:99.0%[S:98.9%,D:0.0%],F:0.3%,M:0.7%,n:6641,E:2.1% |
| CS133 | S_habrochaites | Lima | 2047 | 2 | yes | 61 | 25 | 34958702 | 3122512 | 9 | C:99.0%[S:98.9%,D:0.1%],F:0.3%,M:0.7%,n:6641,E:2.2% |
| CS134 | S_habrochaites | Lima | 2047 | 2 | no | 55 | 23 | 34507928 | 3126575 | 10 | C:99.1%[S:99.0%,D:0.0%],F:0.3%,M:0.7%,n:6641,E:2.1% |
| CS139 | S_lycopersicoides | Arica | 3533 | 1 | no | 30 | 52 | 34726132 | 2644609 | 13 | C:99.0%[S:98.9%,D:0.1%],F:0.3%,M:0.8%,n:6641,E:2.0% |
| CS152 | S_lycopersicoides | Arica | 3533 | 2 | yes | 60 | 30 | 34891182 | 3100262 | 9 | C:98.9%[S:98.9%,D:0.1%],F:0.3%,M:0.8%,n:6641,E:2.0% |

| | | | | | | | | | | | |
|---|---|---|---|---|---|---|---|---|---|---|---|
| CS185 | S_chilense | Tacna | 3091 | 2 | no | 79 | 13 | 34498769 | 3139976 | 9 | C:98.9%[S:98.9%,D:0.1%],F:0.3%,M:0.8%,n:6641,E:2.0% |
| CS311 | S_chilense | San Pedro | 2939 | 1 | no | 39 | 22 | 33955410 | 2872467 | 9 | C:99.0%[S:98.9%,D:0.1%],F:0.3%,M:0.7%,n:6641,E:2.0% |
| CS313 | S_chilense | San Pedro | 2939 | 1 | no | 67 | 58 | 34021790 | 3152621 | 9 | C:99.0%[S:98.9%,D:0.1%],F:0.3%,M:0.8%,n:6641,E:2.0% |
| CS314 | S_chilense | San Pedro | 2939 | 1 | no | 41 | 62 | 34054379 | 2874222 | 10 | C:99.0%[S:98.9%,D:0.1%],F:0.3%,M:0.8%,n:6641,E:2.0% |
| CS330 | S_chilense | Antofagasta | 484 | 0 | yes | 81 | 22 | 34099198 | 3047800 | 9 | C:98.9%[S:98.9%,D:0.0%],F:0.3%,M:0.8%,n:6641,E:2.0% |
| CS339 | S_chilense | Antofagasta | 484 | 1 | no | 39 | 41 | 34376327 | 3095689 | 10 | C:98.9%[S:98.9%,D:0.1%],F:0.3%,M:0.8%,n:6641,E:2.0% |
| CS371 | S_chilense | Arica | 1828 | 1 | no | 14 | 295 | 34987842 | 405082 | 102 | C:98.2%[S:97.7%,D:0.5%],F:0.4%,M:1.4%,n:6641,E:2.1% |
| CS403 | S_chilense | Antofagasta | 75 | 1 | no | 68 | 43 | 34416030 | 2914605 | 11 | C:99.0%[S:98.9%,D:0.1%],F:0.3%,M:0.8%,n:6641,E:2.1% |
| RS002 | S_pimpinellifolium | Lima | 1017 | 1 | no | 42 | 48 | 34142166 | 3085313 | 10 | C:99.0%[S:98.9%,D:0.1%],F:0.3%,M:0.7%,n:6641,E:2.0% |
| RS005 | S_pennellii | Lima | 1017 | 2 | yes | 111 | 15 | 33959535 | 3058231 | 9 | C:99.0%[S:98.9%,D:0.1%],F:0.3%,M:0.7%,n:6641,E:1.9% |
| RS007 | S_pennellii | Lima | 1016 | 1 | no | 57 | 54 | 34571917 | 3189685 | 9 | C:98.9%[S:98.9%,D:0.1%],F:0.3%,M:0.8%,n:6641,E:2.1% |
| RS026 | S_corneliomullerri | Lima | 2900 | 1 | no | 37 | 56 | 34509593 | 2896030 | 10 | C:99.0%[S:98.7%,D:0.3%],F:0.3%,M:0.8%,n:6641,E:2.1% |
| RS040 | S_habrochaites | Lima | 2928 | 1 | no | 59 | 38 | 33589717 | 2906235 | 9 | C:99.0%[S:98.9%,D:0.1%],F:0.3%,M:0.7%,n:6641,E:2.5% |
| RS048 | S_habrochaites | Lima | 1800 | 2 | no | 98 | 16 | 34328983 | 3118230 | 9 | C:99.0%[S:98.9%,D:0.1%],F:0.3%,M:0.8%,n:6641,E:2.1% |
| RS052 | S_habrochaites | Lima | 1900 | 1 | no | 45 | 39 | 34464768 | 3106281 | 10 | C:99.0%[S:98.8%,D:0.2%],F:0.3%,M:0.7%,n:6641,E:1.9% |
| RS071 | S_arcanum | Cajamarca | 2470 | 1 | no | 40 | 31 | 34119171 | 2730057 | 10 | C:99.0%[S:98.9%,D:0.0%],F:0.3%,M:0.7%,n:6641,E:1.9% |
| RS102 | S_habrochaites | Cajamarca | 2085 | 1 | no | 40 | 31 | 34393052 | 3124575 | 10 | C:99.0%[S:98.8%,D:0.2%],F:0.3%,M:0.7%,n:6641,E:2.0% |

Supplementary Table S1 continued

| GC content (%) | Number of annotated genes | Number of CAZYmes | Number of secreted | Number of BGCs | Interspersed repeats (%) | RIP Affected (%) | Count of LRARs | Clade | Mating type | Clone correction | Isolate |
|---|---|---|---|---|---|---|---|---|---|---|---|
| 50,71 | 12622 | 612 | 1196 | 42 | 2,827925467 | 2,51 | 43 | clade1A | MAT1-1 | | CS007 |
| 50,92 | 11952 | 601 | 1161 | 35 | 2,101612356 | 1,78 | 23 | clade1B | MAT1-2 | | CS013 |
| 50,77 | 12005 | 594 | 1159 | 38 | 3,029031389 | 2,53 | 47 | clade1A | MAT1-2 | | CS017 |
| 50,96 | 12023 | 587 | 1131 | 38 | 2,361016558 | 1,97 | 35 | clade2 | MAT1-2 | | CS026 |
| 50,92 | 11768 | 573 | 1086 | 35 | 1,850932507 | 1,84 | 31 | clade1A | MAT1-2 | | CS038 |
| 50,9 | 10708 | 510 | 997 | 30 | 1,930178174 | 1,88 | 25 | clade2 | MAT1-1 | | CS042 |
| 50,97 | 12012 | 591 | 1154 | 36 | 1,923915324 | 1,57 | 25 | clade1B | MAT1-1 | | CS046 |
| 50,96 | 12180 | 600 | 1145 | 38 | 3,065458648 | 1,82 | 33 | clade2 | MAT1-2 | | CS057 |
| 50,91 | 11917 | 586 | 1156 | 37 | 1,828161613 | 1,86 | 28 | clade1B | MAT1-2 | | CS085 |
| 51,08 | 12032 | 589 | 1143 | 35 | 3,318041626 | 1,57 | 32 | clade2 | MAT1-2 | | CS091 |
| 50,99 | 12120 | 596 | 1165 | 37 | 2,03796768 | 1,65 | 20 | clade1C | MAT1-1 | | CS092 |
| 50,96 | 12051 | 595 | 1136 | 36 | 3,040145258 | 1,98 | 34 | clade2 | MAT1-2 | | CS106 |
| 51,01 | 12057 | 599 | 1134 | 37 | 2,753997907 | 1,9 | 42 | clade2 | MAT1-1 | kept | CS126 |
| 51 | 12071 | 602 | 1163 | 37 | 4,0093408 | 1,66 | 31 | clade2 | MAT1-2 | | CS133 |
| 51,03 | 12051 | 591 | 1143 | 37 | 3,31175491 | 1,88 | 41 | clade2 | MAT1-1 | removed | CS134 |
| 50,93 | 12143 | 590 | 1161 | 36 | 2,733920956 | 1,91 | 34 | clade2 | MAT1-1 | | CS139 |
| 51 | 12109 | 585 | 1136 | 35 | 3,108349634 | 1,77 | 38 | clade2 | MAT1-1 | | CS152 |
| 51,01 | 12084 | 594 | 1155 | 38 | 3,092629189 | 1,89 | 32 | clade2 | MAT1-2 | | CS185 |
| 50,93 | 12035 | 588 | 1142 | 35 | 1,844121452 | 1,68 | 25 | clade1C | MAT1-1 | removed | CS311 |
| 50,94 | 12042 | 597 | 1157 | 35 | 1,90270412 | 1,72 | 27 | clade1C | MAT1-1 | kept | CS313 |
| 50,95 | 12002 | 589 | 1150 | 36 | 2,060184389 | 1,72 | 25 | clade1C | MAT1-1 | removed | CS314 |
| 50,91 | 12026 | 593 | 1154 | 36 | 2,369894447 | 1,92 | 34 | clade1A | MAT1-1 | kept | CS330 |
| 50,8 | 12038 | 591 | 1156 | 37 | 2,513406973 | 2,23 | 36 | clade1A | MAT1-1 | removed | CS339 |
| 50,76 | 12077 | 586 | 1113 | 38 | 2,280397859 | 2,37 | 29 | clade1A | MAT1-1 | | CS371 |
| 50,99 | 11967 | 584 | 1154 | 40 | 3,385460206 | 1,87 | 41 | clade2 | MAT1-1 | | CS403 |
| 50,97 | 12023 | 603 | 1164 | 36 | 1,913153372 | 1,62 | 22 | clade1B | MAT1-2 | | RS002 |
| 50,93 | 11944 | 589 | 1146 | 37 | 1,956766691 | 1,74 | 29 | clade1B | MAT1-2 | | RS005 |
| 51,01 | 12079 | 595 | 1142 | 38 | 2,752228058 | 1,58 | 26 | clade2 | MAT1-2 | kept | RS007 |
| 51,03 | 12074 | 597 | 1142 | 38 | 2,763237457 | 1,48 | 25 | clade2 | MAT1-2 | removed | RS026 |
| 50,78 | 11735 | 589 | 1115 | 34 | 2,587553804 | 2,32 | 31 | p.burnsii | MAT1-2 | | RS040 |

| | | | | | | | | | | | |
|---|---|---|---|---|---|---|---|---|---|---|---|
| 51,1 | 12059 | 587 | 1157 | 37 | 3,015163601 | 1,4 | 22 | clade2 | MAT1-1 | | RS048 |
| 50,94 | 12149 | 608 | 1177 | 33 | 2,226328638 | 1,72 | 26 | clade1B | MAT1-2 | | RS052 |
| 50,9 | 12049 | 591 | 1172 | 36 | 2,086290432 | 1,93 | 29 | clade1A | MAT1-1 | | RS071 |
| 50,91 | 12127 | 602 | 1182 | 35 | 1,988980216 | 1,68 | 27 | clade1B | MAT1-1 | | RS102 |

**Supplementary table S2) Population genomic statistics** Based on the SNP data, diversity statistics were calculated with PopGenome. Genetic distances as mean Manhattan distances are based on the distance matrix of the genind_object in adegenet.

| | **all_33samples** | **all_clade1** | **all_clade2** | **\|** | **noclon_28samples** | **noclon_clade1** | **noclon_clade2** |
|---|---|---|---|---|---|---|---|
| **Segregating sites** | 83054 | 42672 | 47939 | \| | 83054 | 42672 | 47939 |
| **Watterson’s Theta** | 20464.24 | 12406.26 | 14743.37 | \| | 21342.65 | 13123.54 | 15448.19 |
| **Nucleotide diversity within** | 13651.70 | 9763.53 | 13127.01 | \| | 14235.69 | 10349.52 | 13340.79 |
| **Tajima's D** | -1.29 | -0.91 | -0.49 | \| | -1.32 | -0.94 | -0.63 |
| **Nucleotide diversity per site** | 0.0074 | 0.0053 | 0.0071 | \| | 0.0077 | 0.0056 | 0.0072 |
| **Number of samples** | 33 | 18 | 15 | \| | 28 | 15 | 13 |
| **-** | - | - | - | - | - | - | - |
| **Mean Manhattan distances** | 616953.5 | 388964.8 | 304651.8 | \| | 612458.8 | 379671.5 | 310543.7 |

### Extended Methods

#### Biological materials

All fungal isolates in this study have been collected from wild tomato plants in Chile and Peru as described in Schmey et al. 2023 [1]. Information about the host plant and sampling region of each sample can also be found in supplementary table S1 above and figure 1 of the main paper. All isolates with a name that starts with RS have been collected in 2018 while sample names with CS have been collected in 2019.

#### DNA extraction and sequencing

To confirm the suitability of Oxford Nanopore Technologies (ONT) sequencing, we sequenced the isolate CS330 on an ONT MinION device. For this, we extracted high molecular weight (HMW) DNA using the protocol described in Einspanier et al. 2022 [2], which is a phenol/chloroform-based method. Library preparation and ONT MinION sequencing were conducted at LMU gene center (Munich, Germany). Additionally, we obtained short read sequencing data for this isolate from the same DNA. Illumina sequencing was performed at the sequencing facility of the technical university of Munich (TUM, Freising-Weihenstephan, Germany). For RNA extraction, the isolate CS330 was grown in liquid culture, on plates with synthetic nutrient-poor agar (SNA) and on plates with potato dextrose agar (PDA). RNA was extracted with a QIAGEN kit and RNA samples of the three growth conditions were pooled before RNA sequencing. The RNA sequencing itself was then performed at the sequencing facility of the technical university of Munich (TUM, Freising-Weihenstephan, Germany).

Next, we performed the same phenol-chloroform DNA extraction on 20 isolates and sequenced them on an ONT PromethION device. Library preparation and PromethION sequencing at the LMU gene center (Munich, Germany) yielded vastly different results, which led to several repetitions of the process and a total of 4 PromethION flowcells being used. DNA for additional short read sequencing of isolate CS046 was extracted using a QIAGEN DNeasy kit and then sequenced by BGI (Hong Kong).

Additional DNA extractions were performed with a KingFisher Flex robot from ThermoFisher Scientific with Mag-Bind Plant DNA DS Kit from QUIAGEN. The DNA of 13 isolates was sequenced on a PromethION flow cell at Senckenberg BiK-F (Frankfurt, Germany). For eight of the 13 additional isolates, short read sequencing data from BGI DNBseq was available (BGI, Hong Kong).

Basecalling was performed by the respective ONT sequence providers. Sequencing yields were assessed with NanoComp (version 1.23.1, [3]). Quality filtering of the sequencing reads was performed using NanoFilt (version 2.8.1, [4]) keeping only reads with a quality better than Q10.

#### Genome assembly and annotation

We used flye (version 2.8.1, [5]) and medaka (version 1.7.1, https://github.com/nanoporetech/medaka ) to assemble the long read data of all samples. Assembly quality was evaluated with quast (version 5.3.0, [6]) and BUSCO (version 5.8.2, [7]). As alternative, we tried the assemblers wtdbg2 (version 2.5, [8]) and different settings of canu (version 2.2, [9]), but they produced more fragmented assemblies with lower BUSCO scores for the majority of samples.

Additional scaffolding and polishing tools like LINKS (version 2.0.1, [10]), SLR [11], Inspector (version 1.0.2, [12]) and more did not improve the assemblies but produced overcorrection artifacts. The samples for which additional short read data was available were polished using three iterations of pilon (version 1.24, [13]) each.

When visualizing assembly contiguity in D-genies dotplots (online, [14]) by mapping our assemblies to well-resolved reference genomes from NCBI (isolates Y784-BC03 ASM2008506v1 and Z7 ASM 1475150v1), two assemblies showed assembly artifacts in the form of artificially fused chromosomes. To manually split these fusions, we mapped our canu assemblies to the flye_medaka assemblies and looked up the coordinates where to split in the paf-file output of minimap2 (version 2.28-r1209, [15]). The superchromosome scaf_1 in the assembly of RS007 was split into scaf_1:1-2837880, scaf_1:2835830-6266848 and scaf_1:6264302-8082830. The superchromosome scaf_3 in the assembly of RS102 was split into scaf_3:1-414269, scaf_3:414270-2769139 and scaf_3:2769140-4849448. Then we used samtools faidx (version 1.19.2, [16]) to extract the sequences of the split scaffolds. We verified the splits in dotplots with the above-mentioned references. All final assemblies were mapped to the reference Y784-BC03 and visualized in D-genies dotplots.

The telomeric identification toolkit tidk (versions 0.2.41 and 0.2.65, [17]) was run with the subcommand search and the expected telomeric repeat sequence TTAGGG and its canonical notation AACCCT. The number of telomeres was then estimated by visual inspection of the output from tidk plot (plots not shown). Additionally, tidk explore was used to confirm that TTAGGG, here reported as AACCCT, is in fact the expected telomeric repeat sequence.

All final assemblies were cleaned and sorted with funannotate clean and funannotate sort on default settings (funannotate version 1.8.17, [18]). We then used Earl Grey (version 4.0.6, [19]) to produce assembly files in which repeats are softmasked. Subsequently, we followed the remaining steps of the funannotate pipeline for annotation, including the integration of AntiSMASH (version 7.1.0, [20]) results.

We used BiG-SCAPE (versions 1.1.5 and version 2.0.0, [21]) to cluster the biosynthetic genes annotated by AntiSMASH. In BiG-SCAPE 2 we performed several runs with different values for –gcf-cutoffs (from 0.3 to 0.8 in steps of 0.05) and compared gene cluster assignment of clusters known to produce the same metabolite. As the known clusters were often split into different families, even when they showed very high synteny, we refrain from mentioning this analysis in the main paper.

#### Identification of putative accessory chromosomes

We generated a concatenated FASTA file containing the genome sequences of all 34 assemblies and nine downloaded reference genomes with less than 100 scaffolds (downloaded from NCBI 6th August 2024). Then we mapped the concatenated fasta on each of the assemblies with minimap2 (version 2.24), followed by conversion to BAM format, sorting, and indexing with samtools (version 1.12). We computed coverage statistics, including mean and median read depth per scaffold, using Mosdepth (version 0.3.3, [22]). Scaffolds were classified as core contigs when their coverage exceeded the mean coverage across all analyzed genomes, while scaffolds greater than 100 kilobase pairs with coverage below the mean were designated as accessory contigs. We validated the classification accuracy in the Integrative Genomics Viewer (IGV v2.12.3, [23]).
For Transposable element (TE) annotation we employed the Extensive De Novo TE Annotator (EDTA v2.2.0, [24]) and generated soft-masked genome files with bedtools (version 2.26.0, [25]). We extracted coding sequences from the annotated GFF3 files (protein annotations from funannotate) using gffread (version 0.12.7, [26]). Finally, we extracted the number of TEs and genes with standard

GNU command-line utilities for density calculation. We calculated gene-wise relative synonymous codon usage (gRSCU) for each gene using the BioKIT package (version0.2.0, [27]).
Statistical analyses were conducted in R v4.3.1 using RStudio v2023.06.0. Wilcoxon rank-sum tests were performed to compare gene-wise RSCU values between core and accessory contigs. Cliff's delta (d) was calculated using the effsize package to quantify the magnitude of differences between groups, with values interpreted as: negligible effect ($|d| < 0.147$), small effect ($0.147 \leqslant |d| < 0.33$), medium effect ($0.33 \leqslant |d| < 0.474$), and large effect ($|d| \geqslant 0.474$).

Several studies test for the presence of the gene for producing the host-specific AAL toxin by conducting a PCR, which we performed as i*n silico* PCR using their primer pairs and the command seqkit amplicon (version 2.8.2, [28]). First, we used the downloaded genome of *A. arborescens* EGS 39-128 (NCBI accession AIIC01000192.1) as positive control. The primer pairs from [29,30] worked with zero mismatches. Then we used the same commands with the same primers on our assemblies. To confirm that the lack of hits is not a matter of primer specificity, we increased the number of possible mismatches stepwise until a sequence was displayed with five mismatches and consequently confirmed with an online blast search against the NCBI nucleotide database that this sequence is not the gene that should have been amplified with the primers.
The gene to synthesize 6-methylsalicylic acid is a different PKS gene that is used as marker for accessory chromosomes of *Alternaria*. We performed the same *in silico* PCR approach as above with the primers for this gene from [31].

#### Phylogeny

We downloaded additional reference genomes from the ncbi website (accession numbers *A. alstroemeriae* ASM3704443v1, *A. atra* ALTATR162, *A. brassica* ASM493672v1, *A. burnsii* ASM1303605v1, *A. consortialis* JCM_1940_assembly_v001, *A. dauci* ASM2550495v1, *A. gaisen* ASM415602v2, *A. gaisen* ASM2234540v1, *A. infectoria* ASM2404317v1, *A. longipes* ASM1905955v1, *A. panax* ASM1970250v1, *A. solani* ASM295215v1, *S. lycopersici* ASM326831v1) and ran BUSCO on them as described above. To generate phylogenies from BUSCO genes, we used the pipeline BUSCO_phylogenomics (https://github.com/jamiemcg/BUSCO_phylogenomics), which produces two outputs. On the one hand, it identifies all BUSCO genes that are single-copy in all samples, aligns them, trims the alignments and then concatenates them into a supermatrix. We used this supermatrix as input for splitstree (version 6.3.30, [32]). Splitstree computes P-distances to generate a Neighbor Net. To increase the number of genes present in the supermatrix for splitstree, we repeated the analysis after excluding the incomplete assembly CS042. On the other hand, the BUSCO_phylogenomics pipeline identifies BUSCO genes that are single-copy in at least four samples and aligns them. After trimming the alignments, a gene tree is generated from each of them, and written to a file of gene trees. We used these gene trees as input for ASTRAL (version 5.7.8, [33]). By computing quartet trees, ASTRAL infers a species tree from the gene trees while accounting for their discordance. We rooted the resulting species tree and plotted it with information about the samples using R 4.4.0 in RStudio (version 2024.04.2+764) with ggtree [34] and further packages ggplot2, ape, dplyr , glue and ggrepel. As the number of input genes was sufficiently high for high confidence, the local posterior probabilities from ASTRAL are shown as support values.

We used Orthofinder (version 2.5.5, [35]) to infer a species tree from the protein sequences listed in the funannotate annotation results. Subsequently, we rooted the tree from Orthofinder using itol [36]. We calculated and visualized a tree comparison in phylo.io [37].

#### k-mer based clustering with mashtree

Mashtree (version 1.4.6, [38]) rapidly compares full genomes following a k-mer strategy. Min-hash distances are calculated between k-mer sketches of the genomes and then used to cluster the genomes into trees. We computed the mashtree with the same samples as the ASTRAL tree and rooted it in itol. Then we compared it to the ASTRAL tree in phylo.io.

Furthermore, we used mashtree on our 33 *A. alternata* genomes (excluding RS040, because it is likely not *A. alternata*) and all available *A. alternata* reference genomes from the NCBI genomes database available at the time (9$^{th}$ January 2025). To this end we downloaded 89 *A. alternata* genomes and 9 *A. tenuissima* genomes. One additional assembly for *A. arborescens* (accession see above) served as possible root. After running mashtree, we saw that two downloaded assemblies had unrealistic branch lengths, likely because they had been mislabelled on the NCBI database, excluded these and ran mashtree again. Finally, we plotted the tree with ggtree in Rstudio.

#### Single Nucleotide Polymorphisms (SNPs)

To investigate single nucleotide polymorphisms (SNPs), we mapped the long-read sequencing data to the reference genome Yref (accession see above) and used the SNP-caller clair3 (version 1.0.10, [39]) as described in [40].
The samples had very different sequencing depths so we applied the maxDP filter on each of the generated g.vcf files with bcftools (version 1.21, [16]) before merging the them. Then we applied further filters with bcftools, namely SnpGap 3 to remove SNPs close to insertions and deletions (INDELs), QUAL 25 to keep only SNPs better than this quality score and DP 10 to keep only SNPs with a coverage of more than ten. Finally, we created a vcf file from all SNPs that pass all filters. For analyses without the sample RS040, which is likely another species, we merged all g.vcf files except RS040 and then performed all subsequent steps in the same fashion.

We used plink (version 1.9, [41]) to perform linkage pruning and perform a principal component analysis (PCA) of the unlinked SNPs, which we then plotted in Rstudio using the R packages tidyverse, grepel, readr and dplyr. Furthermore, we used the linkage pruned results from plink to perform an ADMIXTURE analysis (version 1.3.0, [42]), the results of which we plotted in Rstudio with the packages pophelper, ggplot2 and gridExtra.

We calculated genetic distances between samples as Manhattan distances and plotted a histogram of these pairwise distances in Rstudio with the packages vcfR, poppr, adegenet and ggplot2. We used the histogram and the distance matrix to identify clones. To generate some population genetic statistics, we used Rstudio with the packages dplyr and PopGenome. Data import into PopGenome was facilitated by splitting the vcf file by chromosomes using bcftools. Additionally, we computed the mean Manhattan distances per clade from the distance matrix.

To test for isolation by distance, we employed RStudio. We used bcftools to make a vcf file of only biallelic SNPs and subsequently removed RS040 and the known clones using R to compute the matrix of pairwise genetic distances as manhattan distances. For the geographic distance matrix, we calculated Haversine distances from the coordinates. Then we used a Mantel test with 999 permutations to correlate the two matrices.

#### Pangenome construction

We constructed a pangenome from the 33 *A. alternata* genomes with PanTools (version 4.3.1, [43]) and supplied information about host plants, sampling regions, elevation and affiliation with phylogenetic clades as phenotypes. Then we used the subcommand pantools gene_classification –phenotype to see which genes are specific to the genomes from certain host plants, regions, clades etc. and attempted to identify these specific genes with an online Megablast against the NCBI nucleotide database (https://blast.ncbi.nlm.nih.gov/Blast.cgi).

#### Effector prediction

We identified effectors from our *de novo* assemblies by first extracting proteins predicted to be secreted by funannotate. These were then scored as predicted effectors or not using EffectorP (version v3.0-Fungi, [44]). The predicted proteomes were then organized into predicted orthogroups using OrthoFinder. We classified orthologroups based on whether at least one member was a predicted effector protein. Distribution of orthologous effector groups was visualized with UpSetR [45].

#### Selective sweeps

We used RAiSD (Raised Accuracy in Sweep Detection) (version 4.0, [46]) to detect genomic regions under positive selection. RAiSD identifies selective sweeps based on a composite μ statistic consisting of multiple selection signatures, i.e., reduction in genetic diversity, increased levels of linkage disequilibrium, and shifts in the site frequency spectrum. We did RAiSD separately for each chromosome using a SNP-driven sliding window of 50 SNPs. The windows with the top 0.01% of the μ-statistic distribution were considered putative selective sweeps. The windows were extended by adding 1050 base pairs in both directions of the windows.

We identified the genes located in the window or extended regions with bedtools intersect and the gene annotation produced by funannotate. To identify overrepresented functional categories we performed Gene Ontology (GO) enrichment using the BiNGO plugin (version 3.0.5, within the Cytoscape graphical interface version 3.10.3 [47]).

#### Transposable Elements

To see if repeat content drives the differences in assembly size, we assessed the correlation between assembly size (bp) and repeat content (%) using Pearson's product-moment and Spearman's rank correlation tests in R. We evaluated the normality of each variable with the Shapiro-Wilk test to determine the appropriate correlation method. We performed these analyses on the full dataset as well as on the two clades separately, both before and after removing outliers. The removed outliers were RS040 beause it is not *A. alternata*, but also CS007 from clade 1 because its large genome size could be caused by duplications as seen in the high BUSCO duplicated score and CS042 from clade 2 because the assembly is incomplete and therefore small.

Repeat annotation by EarlGrey, as summarized in the HighLevelCount.summary.txt result, shows the sequence lengths of the repeat classes in each genome assembly. We plotted these results as stacked bar plots of the 33 *A. alternata* samples and the reference genome of *A. alternata* Y784-BC03 with ggplot and dplyr in Rstudio. We sorted the bars by total bar height, i.e., total sequence length of the repeats, and displayed clade 1 with descending and clade 2 with ascending bar heights.

EarlGrey produces TE divergence landscapes. When run on default, the height of the y-axis is determined by the maximum value to be plotted on the y-axis, which means each plot for each assembly has a different y-axis height. We re-plotted the TE divergence landscapes with fixed y-axis height to make them comparable between samples.

To identify segregating starship transposable elements, we scanned the 32 *de novo A. alternata* genomes, as well as the reference isolate Y784-BC03. We used the starfish pipeline [48], retaining elements where the six flanking genes up- and downstream were conserved. We plotted a heatmap of presence and absence of starships in the identified regions sorted by phylogeny with ggtree in RStudio. For region001, we investigated whether the samples with error status also have the starship present by finding the flanking genes in the assemblies with blast-plus. Then we looked at the annotation of the cargo genes that we found between the flanking genes and the contig breaks between the flanking genes.

#### Reproduction

To determine the mating types of our isolates, we conducted a blastn search (blast-plus with Nucleotide-Nucleotide BLAST 2.13.0+, [49]) of the known mating type loci (accession numbers AB00945.1, AB009452.1, AB465670.1, AB444167.1) against our assemblies. To test whether each of the three mating type ratios (overall, clade 1, and clade 2) significantly deviate from a 50:50 distribution, we used binomial tests in Rstudio. Furthermore, we used Pearson's Chi-squared test with Yates' correction to check whether the ratios differ significantly between clades.

The online version of the RIPper [50] computes the occurrence of repeat induced point mutation (RIP) affected regions in sliding windows over the full genome assemblies. Large RIP affected regions (LRARs) and percentage of RIP-affected regions among the genome are reported in the profile.csv results. To determine whether the difference in RIP percentages between our two clades is statistically significant, we used R to run a Welch's t-test, which is a version of the t-test that does not assume equal variance.
